# Accurate Prediction of Antibody Resistance in Clinical HIV-1 Isolates

**DOI:** 10.1101/364828

**Authors:** Reda Rawi, Raghvendra Mall, Chen-Hsiang Shen, Nicole A. Doria-Rose, S. Katie Farney, Andrea Shiakolas, Jing Zhou, Tae-Wook Chun, Rebecca M. Lynch, John R. Mascola, Peter D. Kwong, Gwo-Yu Chuang

## Abstract

Broadly neutralizing antibodies (bNAbs) targeting the HIV-1 envelope glycoprotein (Env) have promising utility in prevention and treatment of HIV-1 infection with several undergoing clinical trials. Due to high sequence diversity and mutation rate of HIV-1, viral isolates are often resistant to particular bNAbs. Resistant strains are commonly identified by time-consuming and expensive *in vitro* neutralization experiments. Here, we developed machine learning-based classifiers that accurately predict resistance of HIV-1 strains to 33 neutralizing antibodies. Notably, our classifiers achieved an overall prediction accuracy of 96% for 212 clinical isolates from patients enrolled in four different clinical trials. Moreover, use of the tree-based machine learning method gradient boosting machine enabled us to identify critical epitope features that distinguish between antibody resistance and sensitivity. The availability of an *in silico* antibody resistance predictor will facilitate informed decisions of antibody usage in clinical settings.

## Introduction

During the past decade, broadly neutralizing antibodies (bNAbs) were isolated from sera of HIV-1 chronically infected donors, with several undergoing clinical trials for use of preventing and treating HIV-1 infection (1–3). As a result of the high sequence diversity and mutation rate of HIV-1, HIV-1 viral isolates can be resistant to a particular bNAb and administration of bNAbs may lead to viral escape (Figure 1a). Resistant viral strains are usually identified by subcloning or synthesizing amplified Envs, producing pseudoviruses, and performing *in vitro* neutralization assays (1), which is time-consuming and expensive.

**Figure 1.**
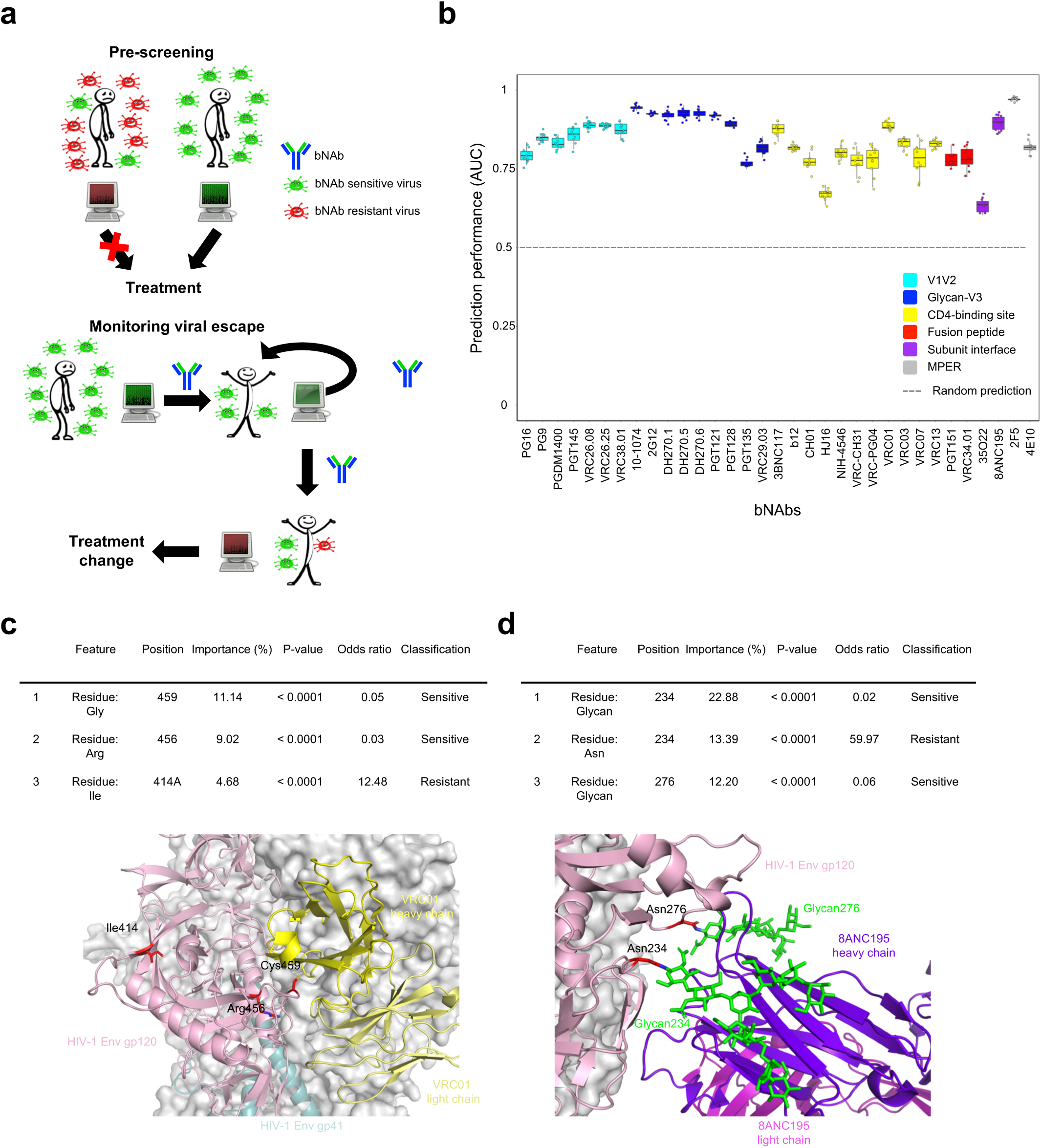
Prediction performance and feature importance of the bNAb-ReP classifiers. **(a)** Schematics illustrating potential applications of bNAb-ReP. **(b)** Prediction performance (AUC) of 33 bNAb classifiers determined by ten runs of ten-fold cross-validation, color-coded based on epitope category, **(c)** The top three discriminant features of the bNAb VRC01 classifier are listed in the table and highlighted on the prefusion-closed Env trimer structure in complex with VRC01 antibody (PDB ID: 5FYJ). **(d)** The top three discriminant features of the bNAb 8ANC195 classifier are listed in the table and highlighted in the Env trimer structure in complex with 8ANC195 bNAb, with glycans 234 and 276 depicted as green sticks (PDB ID: 5CJX).

While many genotypic assays and *in silico* algorithms have been developed to predict HIV-1 drug resistance (4) and co-receptor usage (5), *in silico* prediction of neutralization susceptibility to bNAbs has only been explored by a few studies (6–8), none with publicly available software.

In this work, we present bNAb-Resistance Predictor (bNAb-ReP), an HIV-1 isolate antibody resistance predictor based on the white-box non-linear predictive modeling technique, gradient boosting machine (GBM) (Supplementary Figure S1). GBM has been shown to be competitive with black-box non-linear modeling techniques such as deep learning, particularly when large amounts of training data are not available (9, 10). To build bNAb-ReP, we used sequence and neutralization data for 33 HIV-1 bNAbs obtained from the CATNAP database (11). Full Env sequences and IC_50_ neutralization titers for 205 to 711 HIV-1 isolates with varying clade distributions were available for each bNAb (Supplementary Figure S2).

## Results and discussion

The performance of bNAb-ReP was evaluated in ten runs of ten-fold cross validation measured as the area under the receiver operating characteristic curve (AUC). All bNAb-ReP classifiers performed better than random prediction with average AUC values between 0.63 and 0.97 and an overall median AUC of 0.83 (Figure 1b). Notably, the AUC scores were significantly higher in 26 of the 33 cases using GBM when compared to conventional prediction methods like logistic regression and random forest (Supplementary Figure S3).

In contrast to black-box machine learning approaches, the major advantage of the tree-based method, GBM, is the ability to obtain variable importance scores for all input features, which enables interpretability of the predictive models (feature importance for all 33 bNAb classifiers can be found under https://github.com/RedaRawi/bNAb-ReP). For instance, the top three discriminative features of the bNAb VRC01 classifier involve HIV-1 Env residues 414A, 456, and 459 with a total feature importance of 24.84% (Figure 1c, Supplementary Table S1). Structural studies revealed two of the three amino acid positions were located at the VRC01 epitope and thus can be critical to VRC01 binding and neutralization (Figure 1c) (12). Additionally, the top three features of bNAb 8ANC195 classifier account for a total variable importance of 48.47% and include Env residues 234 and 276, which must be glycosylated in order for 8ANC195 to bind and neutralize Env (Figure 1d, Supplementary Table S2) (13). Not all of the top discriminative features, however, involved bNAb epitope residues, suggesting that distal regions outside the epitope also affect neutralization susceptibility of HIV-1 strains (Supplementary Table S4). In particular, the prediction accuracy for glycan-V3 directed bNAbs had the highest reduction when using only epitope regions rather than full Env sequences. These feature importance scores provide helpful information to facilitate bNAb optimization and guide immunogen design.

To validate bNAb-ReP beyond the data sets obtained from CATNAP, we predicted antibody resistance of HIV-1 Env sequences from clinical studies of HIV infection. First, we tested the bNAb VRC01 classifier on clinical HIV-1 isolates obtained from HIV positive patients enrolled in the VRC601 trial (1) studying the efficacy of VRC01 as therapeutic to control viral load. bNAb-ReP correctly predicted 100% of the resistant and 87% of the sensitive strains to VRC01 (Figure 2a, b).

**Figure 2.**
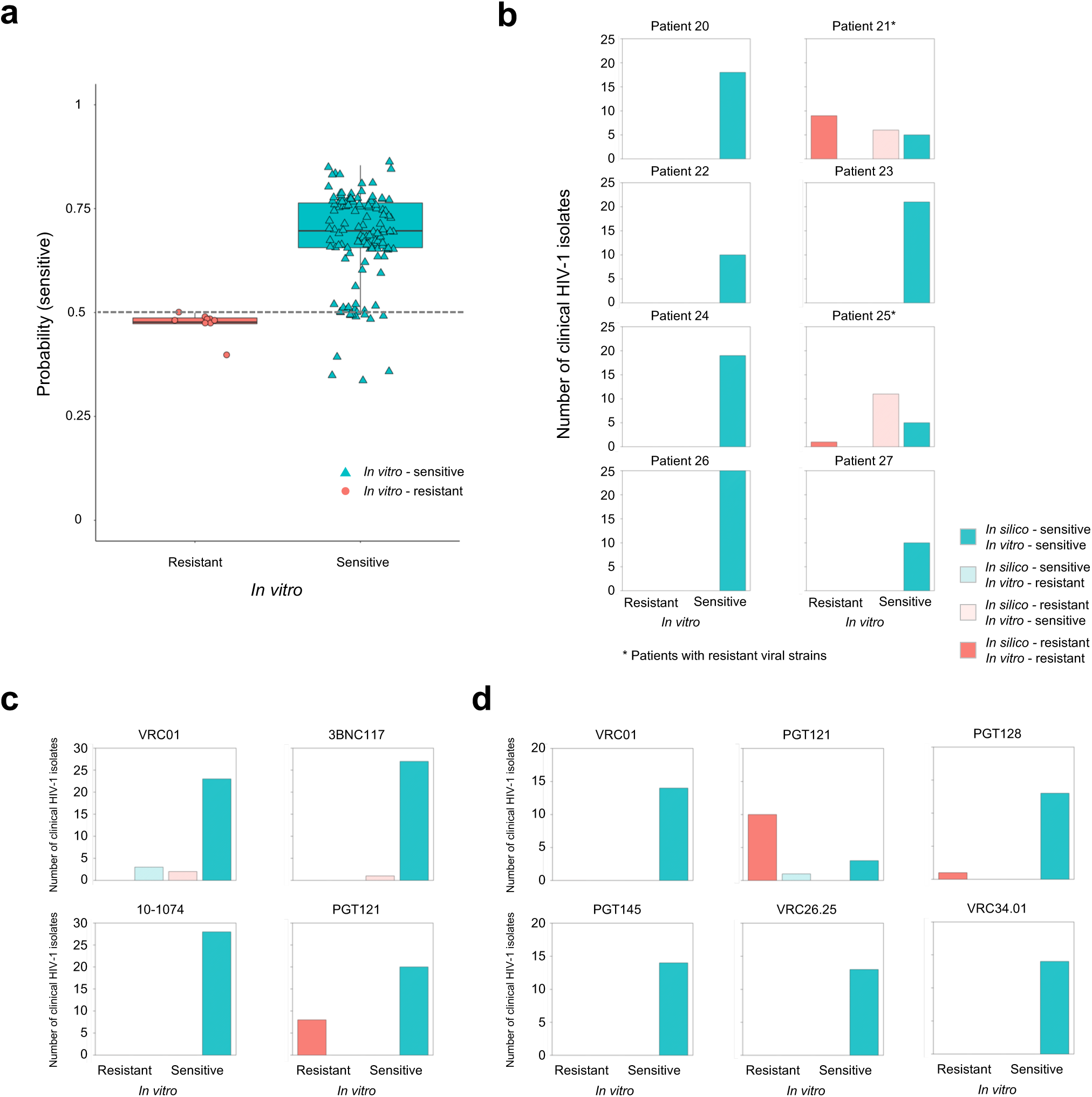
bNAb-ReP prediction performance on clinical HIV-1 isolates. **(a)** Prediction performance of the susceptibility of VRC601 clinical isolates to VRC01. *In vitro* assay neutralization classification is shown on the x-axis, with the *in silico* predicted probability for a sequence to be sensitive to VRC01 shown on the y-axis. The classification cutoff of 0.5 is depicted with a grey dashed line, **(b)** Bar plots depicting the number of *in vitro* classified VRC601 HIV-1 isolates per patient. Clinical HIV-1 isolates *in silico* predictions are shown in red (resistant) and cyan (sensitive) with darker colors indicating true predictions and light colors indicating false predictions.**(c)** Bar plots highlighting the number of clinical HIV-1 isolates, introduced in the Bar et al. study, separated according to their *in silico* predictions. Resistant *in silico* predictions for bNAbs VRC01, 3BNC117, 10-1074, and PGT121 are shown in red and sensitive in cyan, with darker colors representing accurate predictions and light colors inaccurate ones, respectively, **(d)** Bar plots depicting the number of isolates, introduced by Ssemwanga et al., with resistant *in silico* predictions shown in red and sensitive in cyan.

Additionally, we evaluated the prediction performance of bNAb-ReP using sequence and neutralization data from a phase IIa clinical trial studying HIV positive patients treated with bNAb 3BNC117 (3). bNAb-ReP’s overall classification accuracy was 87%, correctly predicting 26 of 29 sensitive HIV-1 Env strains, although falsely predicting the only resistant sequence as sensitive (Supplementary Figure S5).

To further evaluate bNAb-ReP’s prediction accuracy, we performed *in vitro* neutralization assay experiments on clinical sequences obtained from Bar et al. (Supplementary Data S1 and Table S4) (2). bNAb-ReP predicted neutralization susceptibility to VRC01, 3BNC117, 10-1074, and PGT121 with accuracies of 82%, 96%, 100%, and 100%, respectively (Figure 2c).

In addition to predicting neutralization susceptibility from the aforementioned clade B sequences, we used bNAb-ReP to predict resistance for clade A and A/D recombinant sequences from a superinfection case study in a Ugandan couple (14). Interestingly, the bNAb-ReP classification accuracy was 100%, 93%, 100%, 100%, 100%, and 100%, for bNAbs VRC01, PGT121, PGT128, PGT145, VRC26.25, and VRC34.01, respectively (Figure 2e).

In this work, we developed bNAb-ReP, a GBM-based antibody resistance predictor, and demonstrated bNAb-ReP’s ability to predict neutralization susceptibility of HIV-1 isolates with high accuracy for several independent clinical test sets. The underlying machine learning technique, GBM, provided insight into how specific features can distinguish between predicting resistance and sensitivity in HIV-1 strains, informing antibody optimization and immunogen design experiments. The availability of bNAb-ReP will facilitate easy and fast prediction of HIV-1 isolates for their neutralization susceptibility to bNAbs, allowing realtime assessment in a clinical setting. The bNAb-ReP predictors for 33 HIV-1 bNAbs are available for download at https://github.com/RedaRawi/bNAb-ReP.

## Acknowledgements

We thank J. Stuckey and J. Lara for assistance with figures, and members of the Structural Biology Section and Structural Bioinformatics Core, Vaccine Research Center, for discussions or comments on the manuscript. Support for this work was provided by the Intramural Research Program of the Vaccine Research Center, National Institute of Allergy and Infectious Diseases, and by IAVI Neutralizing Antibody Consortium (NAC). This work utilized the computational resources of the NIH HPC Biowulf cluster (NIH HPC). This study used the Office of Cyber Infrastructure and Computational Biology (OCICB) High Performance Computing (HPC) cluster at the National Institute of Allergy and Infectious Diseases (NIAID), Bethesda, MD.

## Author contributions

R.R. and G.-Y.C. designed research; R.R., C.-H. S., R.M., S.K.F., J.Z., performed computational research; N.A.D., A.S., T.-W.C., R.L., and J.R.M. for viral sequence and neutralization data; R.R., P.D.K., and G.-Y.C. wrote the paper, with all authors providing comments or revisions.

## Materials and methods

### Training data

We used neutralization data of 33 different antibodies (10-1074, 2F5, 2G12, 35O22, 3BNC117, 4E10, 8ANC195, CH01, DH270.1, DH270.5, DH270.6, HJ16, NIH-4546, PG16, PG9, PGDM1400, PGT121, PGT128, PGT135, PGT145, PGT151, VRC-CH31, VRC-PG04, VRC01, VRC03, VRC07, VRC13, VRC26.08, VRC26.25, VRC29.03, VRC34.01, VRC38.01 and b12) assayed against 205 to 711 HIV-1 isolates published in the CATNAP database (1). These assays were performed using single-round-of-infection Env-pseudoviruses on cell lines (2, 3). Each HIV-1 isolate is represented with its full-length envelope glycoprotein amino acid sequence. Duplicated full-length HIV-1 envelope sequences were removed. The viral isolate was categorized as resistant to an antibody if its geometric mean IC_50_ is greater than 50μg/ml or designated with a “>” sign or was categorized as sensitive otherwise.

### Test data

For the data from VRC601 clinical trial, we used sequences and neutralization data from Env-pseudoviruses generated from Envs isolated by single genome amplification (SGA) RT-PCR from plasma virus, as described in Lynch et al. (4). The Env-pseudoviruses were assayed on TZM-bl cells as in Sarzotti-Kelsoe et al. (3). The clade A and A/D sequences and neutralization data were generated the same way and were taken from Ssemwanga et al. (5). The VRC01-ATI sequences were derived from patients in an analytical treatment interruption trial, in which volunteers based at the NIH were administered VRC01 infusions before and during an interruption of antiretroviral therapy (6). In that publication, Env sequences were generated by SGA; however, the published neutralization assays were performed with infectious virus from outgrowth cultures, not in the Env-pseudovirus/TZM-bl format. Here, we report new data, for which we expressed the Env-pseudoviruses from the sequences reported in Bar et al. and used the TZM-bl format as above (6).

### Gradient boosting machine (GBM)

To build the training models, we employed a non-linear interpretable tree-based ensemble technique referred as gradient boosting machine (GBM) for building antibody resistance predictors using *h2o* package (Version 3.16.0.2) in *R* software (https://www.R-project.org) (7, 8). GBM belongs to the family of predictive methods, which uses an iterative strategy such that the learning framework will consecutively fit new models so as to have a more accurate estimate of the response variable after each iteration. The primary notion behind this technique is to construct new tree-based learners to be as correlated as possible with the negative gradient of a given loss function, calculated using all the training data. We can use any arbitrary loss function (L(·,·)) here. However, if the loss function is the most commonly used squared-loss function, the learning procedure would result in consecutive residual error-fitting. Algorithm 1 summarizes the generic GBM approach.

#### Algorithm 1: Gradient Boosting Machine

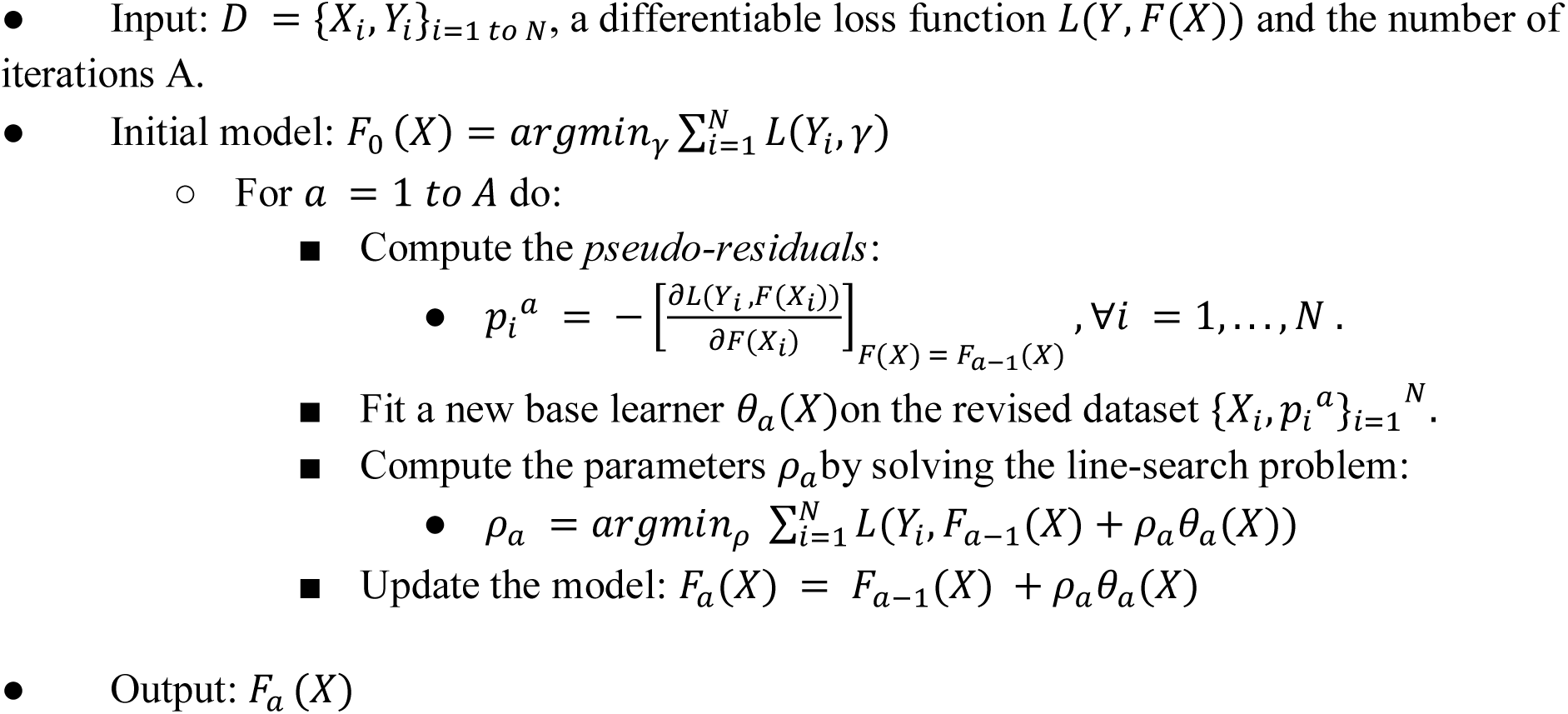

The advantage of the boosting procedure is that it works on decreasing the bias of the model, without increasing the variance. Learning uncorrelated base learners helps to reduce the bias of the final ensemble model. In this work, we used the *L*_2_-TreeBoost approach proposed in (7) to build the core GBM model. Here the loss function is the classical squared-loss function (*L*_2_):

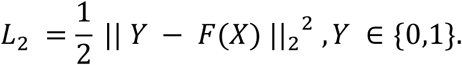

In our approach, the base learner is a *J*-terminal node classification tree. Each tree model has an additive form given as:

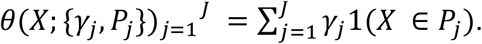

Here 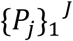 are *J* disjoint regions that together cover the space of all joint values of the predictor variable *X.* These regions represent the J terminal nodes of the corresponding classification tree. The indicator function 1(·) takes the value 1 if the argument passed to it is true, and 0 otherwise. Because the regions are disjoint, *θ*(*X*) is equivalent to the prediction rule: *if X* ∈ *P_j_*, *then θ*(*X*) = *γ_j_*. Now, the pseudo-residuals become:

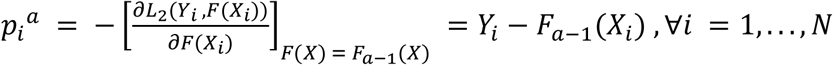

The line search becomes:

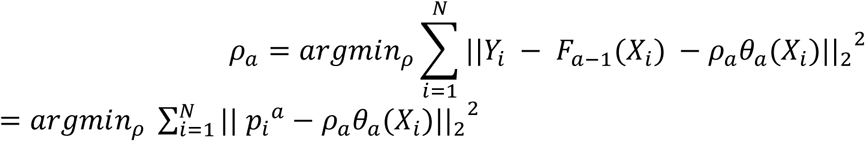

Using classification trees as base learners, we use the idea of separate updates for each terminal region 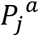 as proposed in (7) to get:

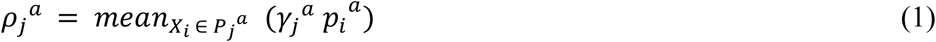

The *L*_2_-TreeBoost approach for two-class GBM is summarized in Algorithm 2.

#### Algorithm 2: *L*_2_-TreeBoost method for GBM

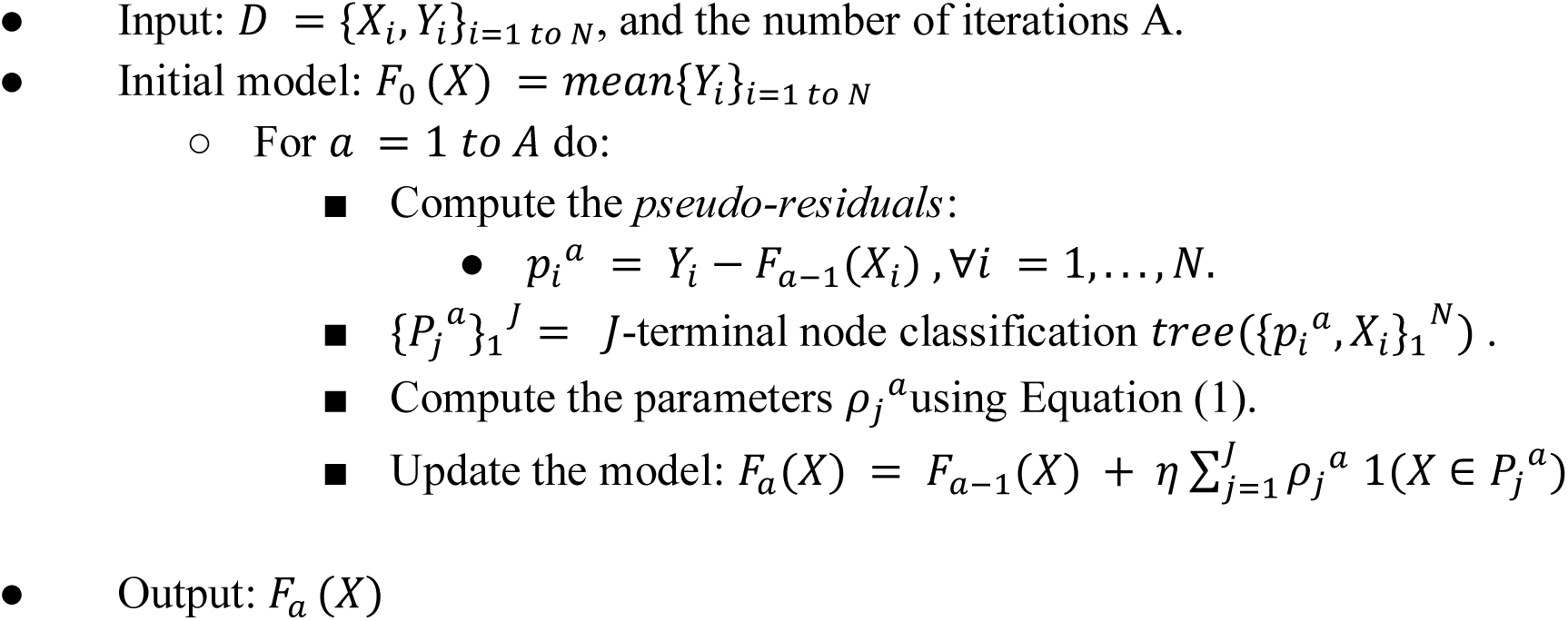

Here the parameter *η* is a regularization parameter which is used to avoid overfitting the models and is acquired via cross validation. For each iteration a, the least-squares criterion (*I*(*ϕ*)) used to assess potential splits of a current terminal region *P* into two disjoint sub-regions (*P_l_*, *P_r_*) is given by:

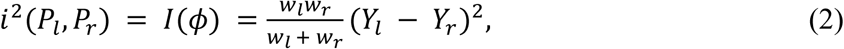

where *Y_l_* and *Y_r_* are the left and right child node responses respectively, and *w_l_ w_r_* are proportional to the number of samples in regions *P_l_* and *P_r_* respectively as show in (Friedman, 2001). *I*(*ϕ*) is a measure of the importance of the variable (*ϕ*) which maximizes this criterion. During a given iteration, only one feature is allowed to cause a split into 2 terminal regions. Thus, in the case of a *J*-terminal node classification tree, we generate *J* – 1 such measures. However, the same feature can generate multiple optimal splits for the *J*-terminal node tree. In such a scenario, we sum the importance of such features to get the total importance of each feature *ϕ* after A iterations. This procedure results in the variable importance scores from the GBM approach.

### Classifier features

Sequence information was represented using one-hot encoding to represent 20 standard amino acids and N-linked glycan. Each amino acid aa_i_, i∈{1, …, 21} was translated into a 21-dimensional vector, where the i^th^ vector position was set to 1, and all other 20 vector positions were set to 0. For instance, applying one-hot encoding to an amino acid sequence of length 100, would be translated into a binary vector of length 2100.

### Training of bNAb-ReP

To train bNAb-ReP classifiers, we first performed a hyperparameter optimization to identify the optimal GBM parameters for the given data. We created a grid of T × J × r × η = 120, in particular number of trees T = 1000, maximum depth J∈{1,2,3,4,5,6}, sample rate 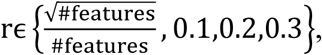 and learn rate η∈{0.001,0.01,0.05,0.1,0.2}. We then performed tenfold cross validation for each of the combinations. Finally, we selected the best parameters that had the maximal ten-fold cross validation area-under-the-curve (AUC). Once the optimal hyper-parameters are known, the model is built on the full training set using these parameters and its prediction performance is evaluated on the independent test set.

### Alternative Predictors Logistic Regression and Random Forest

Logistic Regression belongs to the class of generalized linear models and we trained binomial predictors using *glm* function available in *h2o* package in *R*. Random Forest (RF) belongs to the class of ensemble based supervised white-box learning techniques. The RF algorithm applies the general technique of bagging or bootstrapped aggregating to decision tree learners. We performed a grid search for optimizing the hyper-parameters including the number of trees in the random forest, maximum depth of the trees and column sampling rate using a 10-fold crossvalidation strategy. We used the distributed random forest function, for implementing random forest models, available in *h2o* package in *R*.

### Derivation of probability threshold to categorize sensitivity and resistance

Though there is no clear relationship between the proportion of data to be used for training, testing and the model performance, Shabin et al. identified that the best results were obtained when 75% of the whole dataset was used for training and 25% for testing (9). Similar as implemented by Pfeiffer et al. and Hake et al., we used that probability cutoff as the optimal threshold to distinguish between resistant and sensitive viral sequences (10, 11). In particular, we chose for each bNAb classifier a cutoff that provided the best balance between average true positive and true negative rate.

### Epitope and paratope buried surface area calculations

The buried surface area between antibody and antigen was calculated using NACCESS software (12, 13). The epitope and paratope residues for each antibody were defined as residues with non-zero buried surface area. In the case of 2G12, the epitope residues were defined as glycans N295, N332, N339, N386, and N392, based on Scanlan et al. (14). The final epitope residues for each category were defined as follows. V1V2 category epitope residues comprised all alignment positions between residue numbers 131-196 (HXB2 numbering). The epitope residues for all other categories were defined as the union of all bNAb epitope residues within each category determined as described above.

### Statistical analyses

P-value and odds ratio values presented in Fig.1c and 1d were calculated using Fisher’s exact test (*R* function *fisher.test*). Statistical significance, presented in Supplementary Fig. S3 was determined using the following procedure. First, we tested for the list of AUC values for normal distribution using *R* library *nortest*, in particular function *ad.test.* If normal distribution and additionally variance homogeneity were given (*R* function *var.test*), we used t-test to determine significance (*R* function *t. test*). If neither normal distribution nor variance homogeneity were given, we applied Mann-Whitney test (*R* function *wilcox.test*).

### Data and software availability

We provide all neutralization and sequence data, as well as the resistance predictors for 33 broadly neutralizing HIV-1 antibodies at https://github.com/RedaRawi/bNAb-ReP.

---

**Figure S1.**
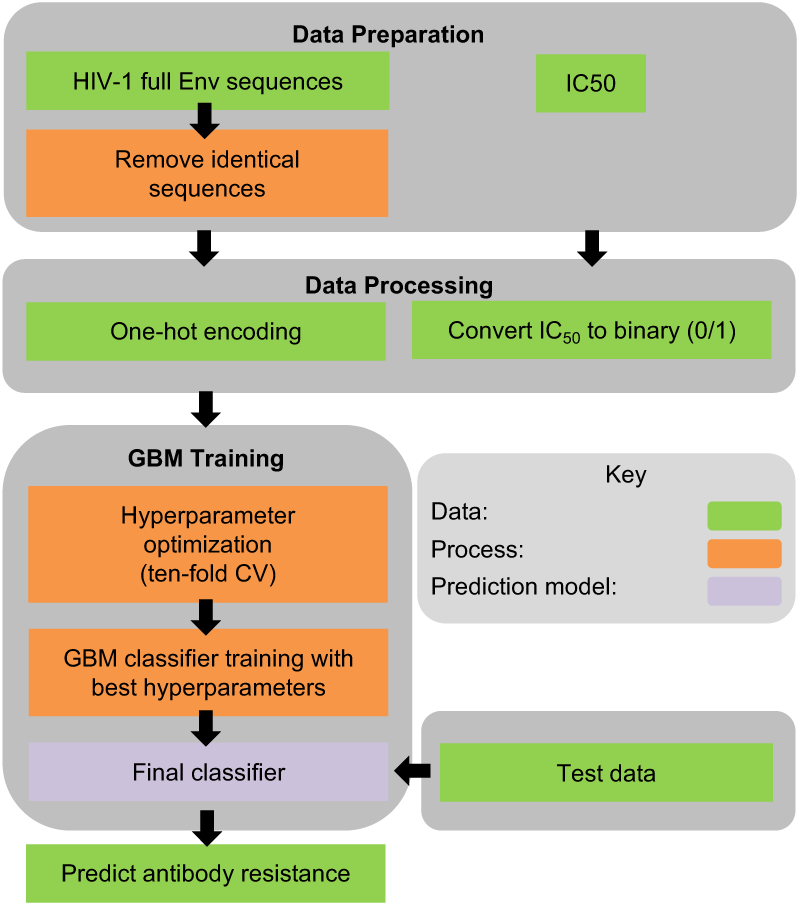
bNAb-ReP development flowchart.

**Figure S2.**
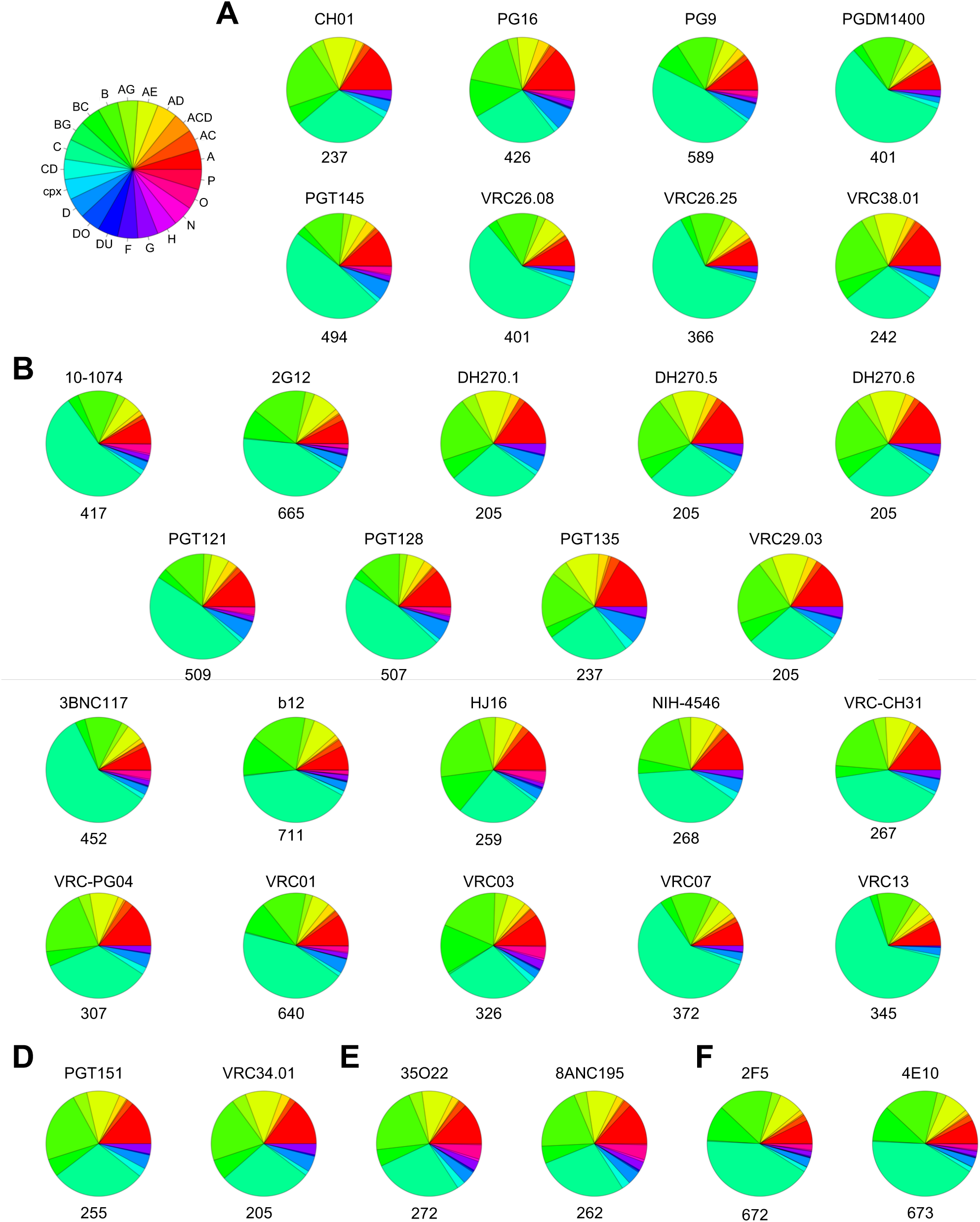
Training sets clade distributions. Clade distributions for each bNAb training set are illustrated as pie charts, with distinct epitope categories shown in (**A**) V1V2, (**B**) Glycan-V3, (**C**) CD4-binding site, (**D**) fusion peptide, (**E**) subunit interface, and (**F**) MPER. The number of strains within each training sets is depicted below the corresponding pie chart.

**Figure S3.**
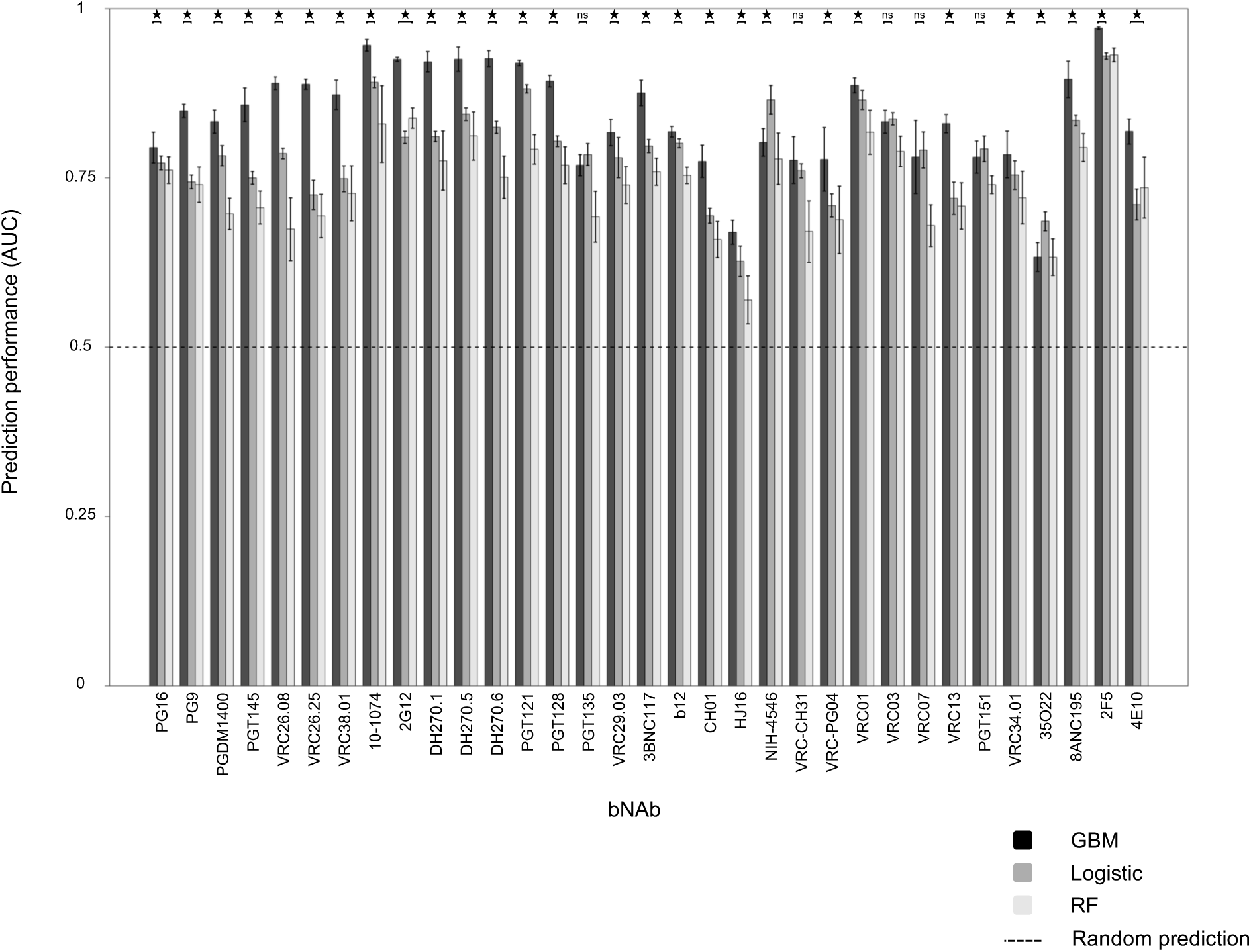
Prediction performance of GBM classifiers compared to Logistic Regression and Random Forest predictors. Prediction performance (AUC) comparison of GBM (black bars), logistic regression (dark gray), and random forest (light gray) of 33 bNAb prediction models determined by ten runs of ten-fold cross-validation. For the sake of clarity, results from statistical significance testing are shown only for the top 2 models (^*^P-value < 0.05, ns = not significant).

**Figure S4.**
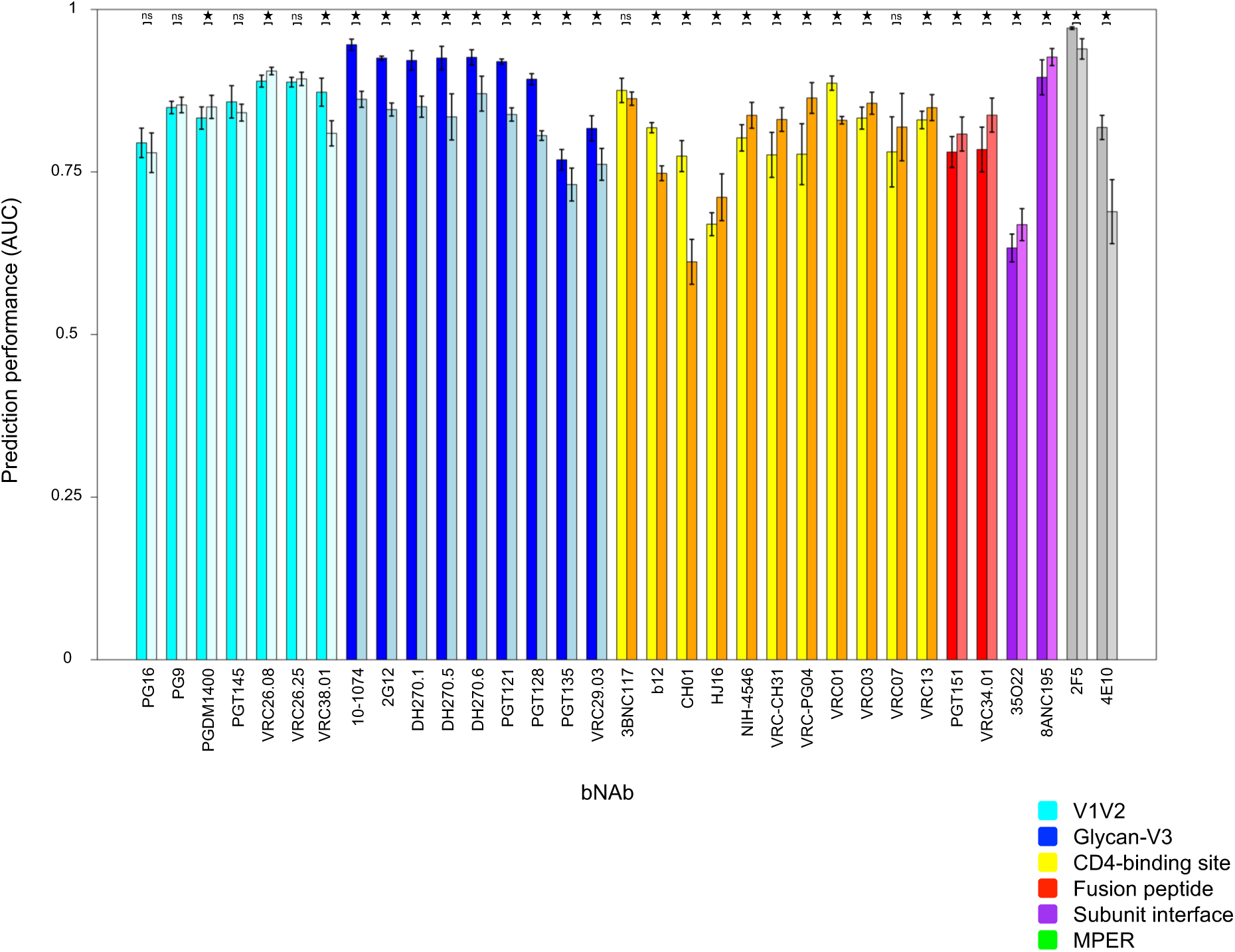
Prediction performance of bNAb classifiers using sequences comprising the full Env sequences or the epitope region only. The prediction performance of the bNAb classifiers using full Env sequences or sequence subsets comprising epitope region only were determined by ten runs of ten-fold cross-validation and are shown as bar plots. The 33 different classifiers are shown on the x-axis, named according to the antibody they are trained on. The colors of the bars refer to the epitope category of the corresponding antibody. The left bar of each pair of bars refers to classifiers trained using full Env sequences, while the right bar refers to classifiers trained using epitope sequence subsets only.

**Figure S5.**
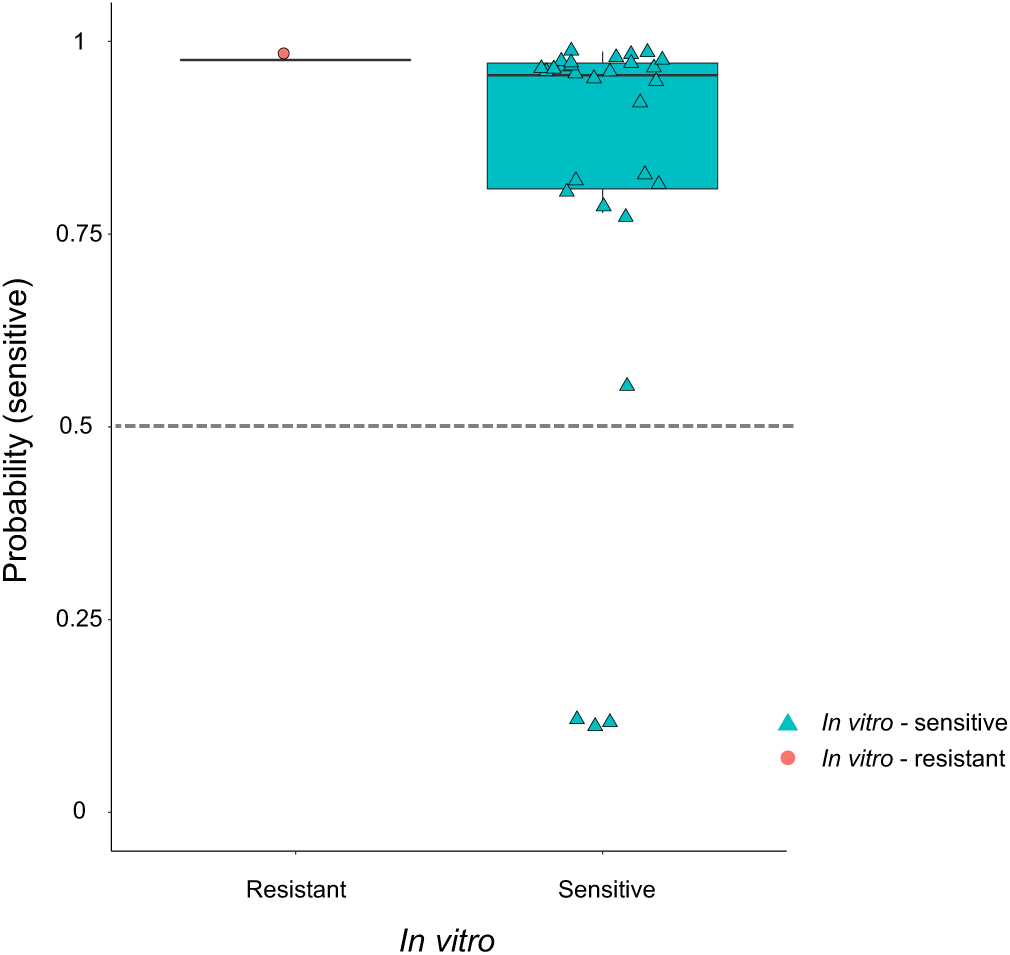
Prediction performance of bNAb-ReP on Env strains from bNAb 3BNC117 treatment study. Prediction performance on bNAb 3BNC117 neutralization and sequence data shown as boxplot. *In vitro* assay neutralization classification is shown on the x-axis, with the *in silico* predicted probability for a sequence to be sensitive to 3BNC117 shown on the y-axis. The classification cutoff of 0.5 is depicted with a grey dashed line.

**Supplementary Table S1.** Feature importance of VRC01 bNAb classifier.

| Feature | Feature importance (%) |
| --- | --- |
| 459_G | 11.13942319 |
| 456_R | 9.020715034 |
| 414A_I | 4.67957789 |
| 365_S | 3.546548512 |
| 280_N | 2.536531598 |
| 364_S | 2.365870823 |
| 588_Q | 2.14043756 |
| 279_D | 2.035677891 |
| 364_H | 1.916603082 |
| 674_S | 1.534500111 |
| 354_glycan | 1.429238443 |
| 839_G | 1.410028298 |
| 106_T | 1.382022626 |
| 269_E | 1.290639271 |
| 396_L | 1.185334014 |
| 274_F | 1.112816848 |
| 602_V | 1.094038602 |
| 413_T | 1.089430625 |
| 853_A | 1.082057062 |
| 389_L | 0.9695362 |
| 50_T | 0.960758772 |
| 400K_glycan | 0.862578797 |
| 335_R | 0.842405863 |
| 720_L | 0.792839621 |
| 655_N | 0.774759909 |
| 399_N | 0.714119092 |
| 804_V | 0.681868304 |
| 340_D | 0.669255526 |
| 639_S | 0.600308988 |
| 458_G | 0.599933891 |
| 316_A | 0.59664073 |
| 362_A | 0.580764573 |
| 740_R | 0.579182655 |
| 534_S | 0.570918655 |
| 723_S | 0.549697325 |
| 340_N | 0.533061329 |
| 525_V | 0.51898338 |
| 856_Q | 0.503857971 |
| 300_S | 0.490333598 |
| 293_S | 0.474870562 |
| 624_G | 0.47324293 |
| 155_R | 0.460370358 |
| 155_K | 0.452165454 |
| 152_K | 0.423181955 |
| 141_V | 0.411366487 |
| 683_R | 0.404268098 |
| 21_I | 0.401669732 |
| 343_Q | 0.392644704 |
| 170_K | 0.384519289 |
| 149_N | 0.377551073 |
| 363_Q | 0.373131642 |
| 192_R | 0.372788451 |
| 49_V | 0.371647651 |
| 851_F | 0.354052754 |
| 703_S | 0.353896049 |
| 444_K | 0.353060438 |
| 403_W | 0.348413332 |
| 730_P | 0.336303169 |
| 348_R | 0.334263924 |
| 429_E | 0.332315171 |
| 316_T | 0.332273048 |
| 275_K | 0.330022143 |
| 279_E | 0.327490564 |
| 139_T | 0.323365064 |
| 417_P | 0.320237403 |
| 276_glycan | 0.317615543 |
| 471_I | 0.314092774 |
| 272_I | 0.310906952 |
| 332_K | 0.310489199 |
| 607_N | 0.306429062 |
| 20_M | 0.291312405 |
| 63_T | 0.288501722 |
| 283_T | 0.288217372 |
| 787D_K | 0.272161993 |
| 151_M | 0.26622777 |
| 279_N | 0.265424596 |
| 154_M | 0.262545237 |
| 16_K | 0.253780916 |
| 812_I | 0.253069063 |
| 335_K | 0.252709111 |
| 114_Q | 0.250375426 |
| 641_T | 0.248055526 |
| 730_L | 0.247050741 |
| 567_K | 0.245379456 |
| 320_T | 0.245378522 |
| 281_A | 0.244095887 |
| 173_F | 0.243904663 |
| 396_N | 0.23971397 |
| 132_G | 0.237710795 |
| 778_A | 0.231182508 |
| 63_R | 0.228742164 |
| 833_G | 0.2238988 |
| 461_glycan | 0.219728075 |
| 333_V | 0.214272549 |
| 209_T | 0.213808467 |
| 80_N | 0.212137819 |
| 4_K | 0.210077905 |
| 144K_T | 0.210043578 |
| 133_glycan | 0.202913049 |
| 86_L | 0.199273638 |
| 463_E | 0.198643696 |
| 334_glycan | 0.195780119 |
| 336_G | 0.191155998 |
| 31_A | 0.190776312 |
| 350_A | 0.190083874 |
| 630_Q | 0.185746992 |
| 130_N | 0.183128064 |
| 31_T | 0.181530723 |
| 535_I | 0.179246405 |
| 492_K | 0.178711947 |
| 138_N | 0.174255959 |
| 400C_W | 0.173352564 |
| 33_K | 0.155484298 |
| 362_T | 0.154288725 |
| 674_D | 0.152903575 |
| 394_glycan | 0.149147735 |
| 141_A | 0.145953436 |
| 375_S | 0.140886931 |
| 373_R | 0.139945948 |
| 519_F | 0.138839296 |
| 769_R | 0.136743099 |
| 169_R | 0.133736679 |
| 412_S | 0.132141771 |
| 798_S | 0.131442345 |
| 186i_S | 0.129983813 |
| 190_E | 0.12972165 |
| 62_D | 0.128477198 |
| 525_A | 0.127908519 |
| 624_D | 0.127032909 |
| 186C_T | 0.126556146 |
| 809_N | 0.125197771 |
| 8_W | 0.124736982 |
| 356_glycan | 0.124495966 |
| 173_S | 0.119938428 |
| 624_N | 0.118608646 |
| 236_T | 0.118136089 |
| 232_K | 0.11357272 |
| 393_K | 0.113443131 |
| 617_K | 0.113246671 |
| 270_I | 0.112952847 |
| 308_S | 0.107290494 |
| 471_G | 0.107203581 |
| 130_glycan | 0.106226507 |
| 144B_glycan | 0.106054614 |
| 702_F | 0.105736095 |
| 143_A | 0.104513555 |
| 4_M | 0.10422366 |
| 411_glycan | 0.10374832 |
| 144J_A | 0.102280728 |
| 85_I | 0.099610797 |
| 4_R | 0.097960213 |
| 800_V | 0.095582969 |
| 641_I | 0.092056504 |
| 746_V | 0.089771612 |
| 297_I | 0.08609011 |
| 352_L | 0.085986627 |
| 389_G | 0.085952406 |
| 50_A | 0.085911514 |
| 460_glycan | 0.08517518 |
| 805_Q | 0.085106445 |
| 616_glycan | 0.084303261 |
| 179_Q | 0.083988168 |
| 171_K | 0.083293309 |
| 106_E | 0.083056525 |
| 352_Y | 0.082507533 |
| 731_G | 0.081395469 |
| 336_D | 0.081366935 |
| 565_L | 0.079195127 |
| 144C_T | 0.075878939 |
| 170_Q | 0.075170581 |
| 446_glycan | 0.074719409 |
| 731_E | 0.074489848 |
| 577_Q | 0.074425197 |
| 416_I | 0.074369649 |
| 156_glycan | 0.074173757 |
| 144_G | 0.074139514 |
| 779_V | 0.074105022 |
| 133_D | 0.073988497 |
| 187_D | 0.071946416 |
| 305_K | 0.071833491 |
| 750_S | 0.071323048 |
| 774_L | 0.071266193 |
| 135_K | 0.071196199 |
| 442_I | 0.07108891 |
| 360_glycan | 0.069722006 |
| 295_V | 0.068672581 |
| 5_G | 0.06858452 |
| 704_V | 0.068197915 |
| 621_D | 0.067989108 |
| 651_N | 0.067464846 |
| 322_D | 0.065793816 |
| 841_C | 0.065441533 |
| 358_K | 0.065424916 |
| 856_L | 0.064983202 |
| 185_N | 0.064409255 |
| 455_V | 0.064171946 |
| 163_T | 0.064161659 |
| 290_E | 0.06246662 |
| 344_K | 0.061986575 |
| 231_E | 0.061908918 |
| 321_G | 0.061648439 |
| 475_M | 0.061066626 |
| 320_N | 0.061035246 |
| 144A_S | 0.060729462 |
| 146_D | 0.060672761 |
| 281_V | 0.060072728 |
| 705_I | 0.058930696 |
| 462_S | 0.058761735 |
| 289_glycan | 0.058516449 |
| 8_P | 0.058481006 |
| 202_A | 0.058272305 |
| 200_A | 0.058100147 |
| 519_L | 0.057872625 |
| 394_T | 0.057777199 |
| 142_V | 0.057625024 |
| 307_I | 0.056932306 |
| 337_Q | 0.056519742 |
| 337_glycan | 0.056450263 |
| 410_P | 0.05643097 |
| 7_Y | 0.056335756 |
| 18_G | 0.055887203 |
| 676_T | 0.055278078 |
| 442_K | 0.054665326 |
| 192_I | 0.054567606 |
| 448_glycan | 0.054007296 |
| 142_G | 0.053999453 |
| 463_glycan | 0.053798537 |
| 186D_S | 0.053071385 |
| 513_I | 0.052428373 |
| 164_S | 0.052265625 |
| 293_Y | 0.05185073 |
| 84_I | 0.051532291 |
| 277_I | 0.050692325 |
| 9_H | 0.050584616 |
| 398_N | 0.050038767 |
| 68_V | 0.049803147 |
| 29_N | 0.048125197 |
| 185_E | 0.04770822 |
| 750_N | 0.047407874 |
| 325_D | 0.047239789 |
| 195_N | 0.046766536 |
| 322_E | 0.046135781 |
| 186D_R | 0.045865955 |
| 270_V | 0.044871106 |
| 668_N | 0.0440979 |
| 607_A | 0.043994047 |
| 293_E | 0.043325789 |
| 753_L | 0.043285088 |
| 770_Q | 0.043234614 |
| 668_S | 0.043207195 |
| 618_T | 0.043007703 |
| 5_E | 0.040770049 |
| 821_A | 0.040640462 |
| 777_G | 0.040533316 |
| 340_E | 0.040279438 |
| 787D_R | 0.039833659 |
| 16_R | 0.039211518 |
| 5B_M | 0.038646184 |
| 211_D | 0.037820515 |
| 756_I | 0.037801195 |
| 732_R | 0.037754135 |
| 136_D | 0.037320003 |
| 33_N | 0.036371353 |
| 690_A | 0.036126926 |
| 658_K | 0.035936344 |
| 169_K | 0.035317706 |
| 209_S | 0.034517785 |
| 63_K | 0.034199306 |
| 363_P | 0.032712572 |
| 351_E | 0.032545602 |
| 395_I | 0.031895506 |
| 754_A | 0.031232184 |
| 467_T | 0.0311497 |
| 619_L | 0.030327124 |
| 291_P | 0.030244051 |
| 440_E | 0.02994854 |
| 165_L | 0.028730217 |
| 158_S | 0.028638126 |
| 179_S | 0.028626081 |
| 395_F | 0.028362373 |
| 354_P | 0.028154713 |
| 60_G | 0.0281164 |
| 496_I | 0.027949231 |
| 173_Y | 0.0279013 |
| 535_V | 0.027720942 |
| 409_S | 0.026881109 |
| 733_I | 0.026607561 |
| 399_G | 0.026596767 |
| 350_R | 0.026417739 |
| 360_E | 0.025972765 |
| 373_T | 0.025533961 |
| 762_S | 0.024962892 |
| 330_H | 0.024360254 |
| 797_G | 0.024260283 |
| 252_R | 0.024114738 |
| 621_E | 0.023685724 |
| 440_A | 0.023498547 |
| 360_Q | 0.023328561 |
| 655_K | 0.023262186 |
| 633_K | 0.022872689 |
| 350_K | 0.022457011 |
| 840_I | 0.021325762 |
| 779_T | 0.021116096 |
| 595_L | 0.021102854 |
| 295_glycan | 0.01945349 |
| 612_S | 0.01936026 |
| 490_E | 0.018880448 |
| 553_S | 0.018762269 |
| 47_E | 0.018735365 |
| 288_L | 0.01823848 |
| 339_glycan | 0.017552792 |
| 779_A | 0.016605813 |
| 831_E | 0.016199103 |
| 275_E | 0.016086297 |
| 459_D | 0.015856236 |
| 413_glycan | 0.015839875 |
| 328_E | 0.015827913 |
| 181_I | 0.01580502 |
| 135_V | 0.015421099 |
| 717_L | 0.015318584 |
| 87_G | 0.015014226 |
| 231_K | 0.014996012 |
| 22_L | 0.014966842 |
| 500_K | 0.014962445 |
| 349_Y | 0.014959779 |
| 695_L | 0.014810424 |
| 494_L | 0.014759353 |
| 300_G | 0.014561998 |
| 809_K | 0.014281161 |
| 290_K | 0.014210631 |
| 184_L | 0.014084515 |
| 21_G | 0.013939846 |
| 364_P | 0.013850563 |
| 782_V | 0.013478092 |
| 758_D | 0.013334677 |
| 128_T | 0.012830445 |
| 63_P | 0.012533033 |
| 85_H | 0.012183287 |
| 613_T | 0.012085381 |
| 46_K | 0.011844576 |
| 240_T | 0.011344416 |
| 362_E | 0.010791987 |
| 5A_I | 0.01073679 |
| 330_Y | 0.010671641 |
| 800_L | 0.010142986 |
| 136_I | 0.009990145 |
| 84_L | 0.009567818 |
| 175_I | 0.009314621 |
| 16_I | 0.009238886 |
| 774_F | 0.009027302 |
| 150_I | 0.008914201 |
| 179_L | 0.008619639 |
| 810_S | 0.008455994 |
| 140_T | 0.008316168 |
| 788_K | 0.008231707 |
| 61_Y | 0.008174783 |
| 161_T | 0.008124686 |
| 767_S | 0.007996554 |
| 388_T | 0.007991136 |
| 142_N | 0.007712182 |
| 84_M | 0.007603355 |
| 85_V | 0.007576291 |
| 354_H | 0.007533432 |
| 46_R | 0.007455503 |
| 742_K | 0.007359883 |
| 19_I | 0.007257145 |
| 459P_T | 0.007206922 |
| 149_S | 0.00714478 |
| 337_K | 0.007117129 |
| 644_S | 0.006813928 |
| 7_C | 0.006812589 |
| 105_Q | 0.006753652 |
| 462_glycan | 0.006726639 |
| 507B_D | 0.006724656 |
| 553_N | 0.006205576 |
| 442_glycan | 0.005506266 |
| 411_S | 0.005368239 |
| 700_A | 0.005367347 |
| 268_E | 0.005346373 |
| 655_R | 0.004987528 |
| 230_glycan | 0.004591001 |
| 153_E | 0.004570718 |
| 662_E | 0.004418435 |
| 186C_glycan | 0.004377783 |
| 429_G | 0.004277492 |
| 295_I | 0.004190532 |
| 99_D | 0.004137473 |
| 762_N | 0.004066218 |
| 25_L | 0.004026744 |
| 171_T | 0.003985033 |
| 717_F | 0.003838424 |
| 345_I | 0.003379397 |
| 60_V | 0.003173227 |
| 132_T | 0.002686051 |
| 29_L | 0.002651904 |
| 787G_Q | 0.00237012 |
| 410_S | 0.002153881 |
| 659_D | 0.002084533 |
| 82_Q | 0.002025609 |
| 344_Q | 0.001658325 |
| 358_glycan | 0.001552825 |
| 281_T | 0.001549052 |
| 300_N | 0.001500181 |
| 784_L | 0.00144775 |
| 240_K | 0.001416826 |
| 741_D | 0.001276793 |
| 149_T | 0.001214967 |
| 839_A | 0.001203856 |
| 26_M | 0.001186211 |
| 136_R | 0.001179731 |
| 776_S | 0.001161185 |
| 587_L | 0.00105952 |
| 659_E | 0.001028506 |
| 283_N | 0.001023159 |
| 414_I | 0.000958059 |
| 700_G | 0.000871272 |
| 61_T | 0.000808637 |
| 287_Q | 0.000768787 |
| 65_V | 0.000671545 |
| 360_V | 0.000658928 |
| 211_E | 0.000557786 |
| 835_R | 0.000457513 |
| 453_I | 0.000428115 |
| 464_N | 0.000426606 |
| 353_F | 0.000404204 |
| 25_I | 0.000385716 |
| 229_K | 0.000357435 |
| 269_K | 0.000349792 |
| 815_L | 0.000348607 |
| 268_G | 0.000320335 |
| 134_V | 0.000319968 |
| 432_Q | 0.000316453 |
| 389_Q | 0.000294976 |
| 92_N | 0.000293384 |
| 15_W | 0.000289998 |
| 325_N | 0.00025003 |
| 565_M | 0.00023286 |
| 842_H | 0.00023235 |
| 698_I | 0.000230161 |
| 154_L | 0.000210466 |
| 408_T | 0.000206756 |
| 99_N | 0.000201348 |
| 4_T | 0.000180081 |
| 821_V | 7.34E-05 |
| 336_T | 6.35E-05 |
| 261_L | 4.76E-05 |
| 322_G | 4.21E-05 |
| 813_N | 3.48E-05 |
| 851_L | 2.35E-05 |
| 332_glycan | 2.22E-05 |
| 395_Y | 1.11E-05 |
| 386_N | 4.17E-06 |

**Supplementary Table S2.** Feature importance of 8ANC195 bNAb classifier.

| Feature | Feature importance (%) |
| --- | --- |
| 234_glycan | 22.88495943 |
| 234_N | 13.39213185 |
| 276_glycan | 12.19668815 |
| 349_L | 7.762024202 |
| 236_T | 4.778803674 |
| 15_W | 4.39447887 |
| 236_K | 3.054029751 |
| 278_T | 2.653325309 |
| 567_K | 2.460009546 |
| 272_V | 2.40779405 |
| 443_I | 1.971399969 |
| 513_I | 1.7595767 |
| 632_E | 1.666828959 |
| 8_Q | 1.426869548 |
| 230_glycan | 1.212115917 |
| 346_T | 1.174123618 |
| 440_A | 1.129961841 |
| 336_A | 1.10929718 |
| 685_F | 1.000928614 |
| 405_S | 0.959278137 |
| 271_K | 0.726755291 |
| 743_D | 0.697478472 |
| 151_G | 0.696037803 |
| 496_V | 0.675314213 |
| 166_R | 0.602167627 |
| 348_K | 0.582416156 |
| 85_F | 0.531671648 |
| 16_R | 0.506053363 |
| 667_K | 0.50392464 |
| 240_N | 0.492548113 |
| 732_G | 0.401376607 |
| 393_S | 0.38705244 |
| 364_P | 0.347079916 |
| 340_K | 0.319151132 |
| 283_T | 0.285876793 |
| 5B_Q | 0.270768944 |
| 440_Q | 0.269014747 |
| 46_K | 0.26586422 |
| 630_Q | 0.26541092 |
| 240_T | 0.240219195 |
| 851_L | 0.220587454 |
| 842_N | 0.168745735 |
| 740_R | 0.147150987 |
| 430_A | 0.145863734 |
| 651_N | 0.136247106 |
| 68_V | 0.121859449 |
| 491_I | 0.1027721 |
| 271_V | 0.096919688 |
| 496_L | 0.085356495 |
| 726_G | 0.07429487 |
| 234_S | 0.051492766 |
| 398_glycan | 0.037094026 |
| 734_G | 0.032201725 |
| 268_G | 0.029990807 |
| 636_N | 0.023519764 |
| 724_P | 0.020782199 |
| 818_T | 0.014454949 |
| 20_M | 0.0142045 |
| 632_D | 0.006849514 |
| 677_K | 0.005064027 |
| 442_glycan | 0.001793042 |
| 837_C | 0.001338458 |
| 182_V | 0.000363412 |
| 787C_L | 0.000245629 |

**Supplementary Table S3.** Overlap of residue position with a feature importance of at least 5% (structurally determined epitope positions are highlighted in red).

| bNAb | Number of features | Number of features (epitope residues) | Residue positions |
| --- | --- | --- | --- |
| b12 | 1 | 0 | 185 |
| 4E10 | 2 | 1 | 674, 787B |
| 2F5 | 4 | 3 | 492, 665, 665, 667 |
| 2G12 | 3 | 2 | 295, 332, 395 |
| VRC01 | 2 | 2 | 456, 459 |
| PG9 | 2 | 2 | 160, 169 |
| PGT128 | 3 | 1 | 332, 334, 334 |
| PGT145 | 2 | 2 | 160, 169 |
| 3BNC117 | 4 | 2 | 456, 459, 466, 723 |
| PG16 | 1 | 1 | 160 |
| 10-1074 | 3 | 1 | 332, 334, 334 |
| VRC13 | 3 | 2 | 179, 471, 471 |
| VRC03 | 0 | 0 |  |
| VRC-PG04 | 7 | 7 | 276, 364, 365, 389, 429, 456, 459 |
| 35O22 | 0 | 0 |  |
| NIH45-46 | 3 | 2 | 279, 364, 456 |
| VRC-CH31 | 2 | 2 | 276, 459 |
| 8ANC195 | 4 | 3 | 234, 234, 276, 349 |
| HJ16 | 1 | 1 | 471 |
| PGT151 | 5 | 2 | 514, 519, 602, 629, 651 |
| VRC38.01 | 2 | 2 | 130, 171 |
| PGT135 | 4 | 1 | 179, 332, 334, 592 |
| VRC34.01 | 1 | 1 | 518 |

**Supplementary Table S4.**
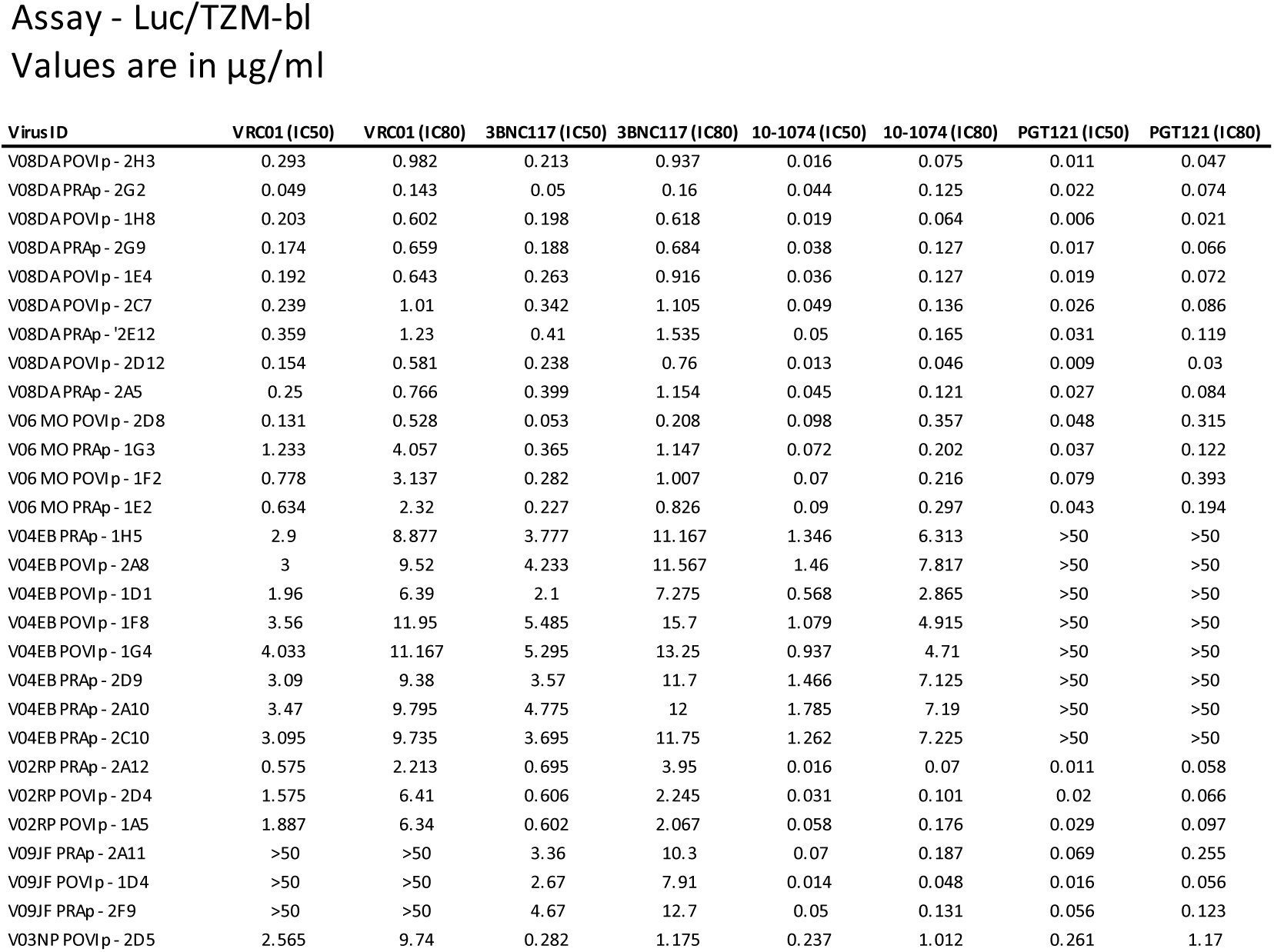
Neutralization data of VRC01-ATI study.

## Supplementary Data S1. HIV-1 Env sequences of VRC01-ATI study

>V08DA_POVIp_2H3.A.9

MRVRGIRKNYQRLWKWG-TLLSMLLGLLMICNAKEQLWVTVYYGVPVWKEANTTLFCASDARAYSTEKHNVWATHACVPTDPDPQEVKLGNVTENFNMWKNNMVDQMHEDIISLWDQSLKPCVEITPLCVTLNCTDYKGNASSTN------NSGGEMENKEEMKNCSFNITTSLSDKV-QERALFYIYDITPINN-----TTNTSYKLTRCNSSVITQACPKVSFEPIPIHYCAPAGFAILKCRQKNFKGTGPCTDVSTVQCTHGIRPVVSTQLLLNGSLAEDEVVIKSENLTDNSKNIIVQLQEAINITCTRPNNNTRKSINI--GPGRAIYATGEIIGDIRQAHCNISEEKWNKTLGQIVEKLRKQFN-NKTIVFAQPSGGDPEIVMHSFNCGGEFFYCNTTRLFNSTWSTWNDS-STLSNGTGNNTITLPCRIKQVINMWQKVGKAMYAPPIAGQIRCSSNITGLILTRDGG----SNNMSNETFRPGGGNMKDNWRSELYKYKVVKIEPLGIAPTKAKRRVVQREKRAIG-IGAVFLGFLGAAGSTMGAASVTLTVQARQLLFGIVQQQNNLLRAIEAQQHLLQLTVWGIKQLQARVLAVERYLKDQQLLGIWGCSGKLICTTAVPWNTSWSNKSLEQIWDNMTWMQWEKEINNYTGLIYTLIEQSQNQQEKNEQELLELDTWASLWNWFDISKWLWYIKIFIMIVGGLIGLRIVFTVLSIVNRVRQGYSPLSFQTRLPAQRG-PDRPEGIEEEGGERDRDRSGPLVDGFFAIIWVDLRNLCLFIYHRLRDLLLIVTRIVELLGR-RGWEILKYWWNLLQYWRQELKNSAISLLNATAIAVAEGTDRVIEVVQRIGRGILHIPVRIRQGLERALL

>V08_DA_PRAp_2G2.2

MRVRGIRKNYQRLWKWG-TLLSMLLGLLMICNAKEQLWVTVYYGVPVWREANTTLFCASDARAYSTEKHNVWATHACVPTDPDPQEVKLGNVTENFNMWKNNMVDQMHEDIISLWDQSLKPCVEITPLCVTLNCTDYRGNDNSTN------NSNETIVNKEEIKNCSFNITTSLSDKR-QERALFYKYDITPINS-----TTNTSYRLTRCNSSVITQACPKVSFEPIPIHYCAPAGFAILKCRQKNFKGTGPCTDVSTVQCTHGIRPVVSTQLLLNGSLAEEEVVIKSESFTDNSKNIIVQLKEAINITCTRPNNNTRKSINI--GPGRAIYATGEIIGDIRQAHCNISEEKWNETLGQIVKKLQEQFG-NKSIIFAQPAGGDPETVMHSFNCGGEFFYCNTTRLFNSTWNDSS----TRSNGTGNNTITLPCRIKQVINMWQKVGKAMYAPPIAGQIRCSSNITGLILTRDGG----SNNMSNETFRPGGGNMKDNWRSELYKYKVVKIEPLGIAPTKAKRRVVQREKRAIG-IGAVFLGFLGAAGSTMGAASVTLTVQARQLLFGIVQQQNNLLRAIEAQQHLLQLTVWGIKQLQARVLAVERYLKDQQLLGIWGCSGKLICTTAVPWNISWSNKSLEQIWNNMTWMQWEKEINNYTGLIYTLIEQSQNQQEKNEQELLELDTWASLWNWFDISKWLWYIKIFIMIVGGLIGLRIVFTVLSIVNRVRQGYSPLSFQTRPPAQRG-PDRPEGIEEEGGERDRDRSGPLVDGFFAIIWVDLRNLCLFIYHRLRDLLLIVTRIVELLGR-RGWEILKYWWNLLQYWRQELKNSAISLLNATAIAVAEGTDRVIEVVQRIGRGILHIPVRIRQGLERALL

>V08DA_POVIp_1H8.6

MRVRGIRKNYQRLWKWG-TLLSMLLGLLMICNAKEQLWVTVYYGVPVWKEANTTLFCASDARAYSTEKHNVWATHACVPTDPDPQEVGLGNVTENFNMWKNNMVDQMHEDIISLWDQSLKPCVEITPLCVTLNCTDYRGNDNSTN------NSNETIVNKEGIKNCSFNITTSLSDKR-QERALFYKYDITPINN-----TTNTSYKLTRCNSSVITQACPKVSFEPIPIHYCAPAGFAILKCRQKNFNGTGPCTNVSTVQCTHGIRPVVSTQLLLNGSLAEEEVVIKSENFTDNSKNIIVQLKEAINITCTRPNNNTRKSINI--GPGRAIYATGEIIGDIRQAHCNISEEKWNETLGQIVEKLQKQFG-NKSIIFAQPSGGDPETIMHSFNCGGEFFYCNTTQLFNSTWNDSS----TRSNGTGNNTITLPCRIKQVINMWQKVGKAMYAPPIAGQIRCSSNITGLILTRDGG-----SNMSNETFRPGGGNMKDNWRSELYKYKVVKIEPLGIAPTKAKRRVVQREKRAIG-IGAVFLGFLGAAGSTMGAASVTLTVQARQLLFGIVQQQNNLLRAIEAQQHLLQLTVWGIKQLQARVLAVERYLKDQQLLGIWGCSGKLICTTAVPWNISWSDKSLEQIWNNMTWMQWEKEINNYTGLIYTLIEQSQNQQEKNEQELLELDTWASLWNWFDISKWLWYIKIFIMIVGGLIGLRIVFTVLSIVNRVRQGYSPLSFQTRPPAQRG-PDRPEGIEEEGGERDRDRSGPLVDGFFAIIWVDLRNLCLFIYHRLRDLLLIVTRIVELLGR-RGWEILKYWWNLLQYWRQELKNSAISLLNATAIAVAEGTDRVIEVVQRIGRGILHIPVRIRQGLERALL

>V08_DA_PRAp_2G9.7

MRVRGIRKNYQRLWKWG-TLLSMLLGLLMICNAKEQLWVTVYYGVPVWEEANTTLFCASDARAYSTEKHNVWATHACVPTDPDPQEVGLGNVTENFNMWKNNMVDQMHEDIISLWDQSLKPCVEITPLCVTLNCTDYRGNDNSTN------NSNETIVNKEEIKNCSFNITTSLSDKV-QERALFYKYDITPINS-----TTNTSYKLTRCNSSVITQACPKVSFEPIPIHYCAPAGFAILKCRQKNFNGTGPCTEVSTVQCTHGIRPVVSTQLLLNGSLAEGEVVIKSENFTDNSKNIIVQLKEAINITCTRPNNNTRKSINI--GPGRAIYATGEIIGDIRQAHCNISEEKWNETLGQIVEKLQEQFG-NKSIIFTQPSGGDPEIVMHSFNCGGEFFYCNTTRLFNSTWNDSS----TRSNGTGNNTITLPCRIKQVINMWQKVGKAMYAPPIAGQIRCSSNITGLILTRDGG----SNNMSNETFRPGGGNMKDNWRSELYKYKVVKIEPLGIAPTKAKRRVVQREKRAIG-IGAVFLGFLGAAGSTMGAASVTLTVQARQLLFGIVQQQNNLLRAIEAQQHLLQLTVWGIKQLQARVLAVERYLKDQQLLGIWGCSGKLICTTAVPWNASWSNKSLEQIWNNMTWMQWEKEINNYTGLIYTLIEQSQNQQEKNEQELLELDTWASLWNWFDISKWLWYIKIFIMIVGGLIGLRIVFTVLSIVNRVRQGYSPLSFQTRPPAQRG-PDRPEGIEEEGGERDRDRSGPLVDGFFAIIWVDLRNLCLFIYHRLRDLLLIVTRIVELLGR-RGWEILKYWWNLLQYWRQELKNSAISLLNATAIAVAEGTDRVIEVVQRIGRGILHIPVRIRQGLERALL

>V08DA_POVIp_1E4.1

MRVRGIRKNYQRLWKWG-TLLSMLLGLLMICNAKEQLWVTVYYGVPVWKEASTTLFCASDARAYSTEKHNVWATHACVPTDPDPQEVGLGNVTENFNMWKNNMVDQMHEDIISLWDQSLKPCVEITPLCVTLNCTDYRGNDNSTN------NSNETIANKEEIKNCSFNITTSLSDKV-QERALFYMYDITPINN-----TTNTSYKLTRCNSSVITQACPKVSFEPIPIHYCAPAGFAILKCRQKNFNGTGPCTNVSTVQCTHGIRPVVSTQLLLNGSLAEEEVVIKSENITDNSKNIIVQLKEAINITCTRPNNNTRKSINI--GPGRAIYATGEIIGDIRQAHCNISEEKWNETLRQIVEKLQKQFG-NKSIIFAQPAGGDPETVMHSFNCGGEFFYCNTTRLFNSTWSTWNDS-STLSNGTGNNTITLPCRIKQVINMWQKVGKAMYAPPIAGQIRCSSNITGLILTRDGG----SNNMSNETFRPGGGNMKDNWRSELYKYKVVKIEPLGIAPTKAKRRVVQREKRAIG-IGAVFLGFLGAAGSTMGAASVTLTVQARQLLFGIVQQQNNLLRAIEAQQHLLQLTVWGIKQLQARVLAVERYLKDQQLMGIWGCSGKLICTTAVPWNASWSNKSLQQIWDNMTWMQWEKEINNYTGLIYTLIEQSQNQQEKNEQELLELDTWASLWNWFDISKWLWYIKIFIMIVGGLIGLRIVFTVLSIVNRVRQGYSPLSFQTRPPAQRG-PDRPEGIEEEGGERDRDRSGPLVDGFFAIIWVDLRNLCLFIYHRLRDLLLIVTRIVELLGR-RGWEILKYWWNLLQYWRQELKNSAISLLNATAIAVAEGTDRVIEVVQRIGRGILHIPVRIRQGLERALL

>V08DA_POVIp_2C7.1

MRVRGIRKNYQRLWKWG-TLLSMLLGLLMICNAKEQLWVTVYYGVPVWKEANTTLFCASDARAYSTEKHNVWATHACVPTDPDPQEVGLGNVTENFNMWKNNMVDQMHEDIISLWDQSLKPCVEITPLCVTLNCTDYKGNDNSTN------NSNETIVNKEEIKNCSFNITTSLSDKV-QERALFYMYDITPINN-----TTNTSYRLTRCNSSVITQACPKVSFEPIPIHYCAPAGFAILKCRQKNFNGTGPCTNVSTVQCTHGIRPVVSTQLLLNGSLAEEEVVIKSENFTDNSKNIIVQLKEAINITCTRPNNNTRKSINI--GPGRAIYATGEIIGDIRQAHCNISEGKWNKTLGQIVEKLQKQFG-NKSIIFTQPSGGDPEIVMHSFNCGGEFFYCNTTRLFNSTWNDSS----TRSNGTGNNTITLPCRIKQVINMWQKVGKAMYAPPIAGQIRCSSNITGLILTRDGG----SNNMSNETFRPGGGNMKDNWRSELYKYKVVKIEPLGIAPTKAKRRVVQREKRAIG-IGAVFLGFLGAAGSTMGAASVTLTVQARQLLFGIVQQQNNLLRAIEAQQHLLQLTVWGIKQLQARVLAVERYLKDQQLLGIWGCSGKLICTTAVPWNTSWSDKSLEQIWNNMTWMQWEKEINNYTGLIYTLIEQSQNQQEKNEQELLELDTWASLWNWFDISKWLWYIKIFIMIVGGLIGLRIVFTVLSIVNRVRQGYSPLSFQTRPPAQRG-PDRPEGIEEEGGERDRDRSGPLVDGFFAIIWVDLRNLCLFIYHRLRDLLLIVTRIVELLGR-RGWEILKYWWNLLQYWRQELKNSAISLLNATAIAVAEGTDRVIEVVQRIGRGILHIPVRIRQGLERALL

>V08_DA_PRAp_2E12.5

MRVRGIRKNYQRLWKWG-TLLSMLLGLLMICNAKEQLWVTVYYGVPVWKEANTTLFCASDARAYSTEKHNVWATHACVPTDPDPQEVGLGNVTENFNMWKNNMVDQMHEDIISLWDQSLKPCVEITPLCVTLNCTDYRGNDNSTN------NSNETIVNKEEIKNCSFNITTSLSDKV-QERALFYMYDITPINN-----TTNTSYKLTRCNSSVITQACPKVSFEPIPIHYCAPAGFAILKCRQKNFNGTGPCTDVSTVQCTHGIRPVVSTQLLLNGSLAEEEVVIKSENFTDNSKNIIVQLKEAINITCTRPNNNTRKSINI--GPGRAIYATGEIIGDIRQAHCNISEEKWNKTLGQIVKKLQKQFG-NKSIIFAQPSGGDPEIVMHSFNCGGEFFYCNTTRLFNSTWSTWNDS-STGSNGTGNNTITLPCRIKQVINMWQKVGKAMYAPPIAGQIRCSSNITGLILTRDGG----SNNMSNETFRPGGGNMKDNWRSELYKYKVVKIEPLGIAPTKAKRRVVQREKRAIG-IGAVFLGFLGAAGSTMGAASVTLTVQARQLLFGIVQQQNNLLRAIEAQQHLLQLTVWGIKQLQARVLAVERYLKDQQLLGIWGCSGKLICTTAVPWNTSWSNKSLQQIWDNMTWMQWEKEINNYTGLIYTLIEQSQNQQEKNEQELLELDTWASLWNWFDISKWLWYIKIFIMIVGGLIGLRIVFTVLSIVNRVRQGYSPLSFQTRPPAQRG-PDRPEGIEEEGGERDRDRSGPLVDGFFAIIWVDLRNLCLFIYHRLRDLLLIVTRIVELLGR-RGWEILKYWWNLLQYWRQELKNSAISLLNATAIAVAEGTDRVIEVVQRIGRGILHIPVRIRQGLERALL

>V08DA_POVIp_2D12.A.3

MRVRGIRKNYQRWWKWG-TLLSMLLGLLMICNAKEQLWVTVYYGVPVWKEASTTLFCASDARAYSTEKHNVWATHACVPTDPDPQEVGLGNVTENFNMWKNNMVDQMHEDIISLWDQSLKPCVEITPLCVTLNCTDYRGND----------NSNETIVNKEEIKNCSFNITTSLSDKV-QERALFYMYDITPINN-----TTNTSYKLTRCNSSVITQACPKVSFEPIPIHYCAPAGFAILKCRQKNFNGTGPCTDVSTVQCTHGIRPVVSTQLLLNGSLAEGEVVIKSENFTDNSKNIIVQLKEAINITCTRPNNNTRKSINI--GPGRAIYATGEIIGDIRQAHCNISEEKWNKTLGQIVEKLQKQFG-NKSIIFAQPSGGDPETVMHSFNCGGEFFYCNTTRLFNSTWNDSS----TLSNGTGNNTITLPCRIKQVINMWQKVGKAMYAPPIAGQIRCSSNITGLILTRDGG----SNNMSNETFRPGGGNMKDNWRSELYKYKVVKIEPLGIAPTKAKRRVVQREKRAIG-IGAVFLGFLGAAGSTMGAASVTLTVQARQLLFGIVQQQNNLLRAIEAQQHLLQLTVWGIKQLQARVLAVERYLKDQQLLGIWGCSGKLICTTAVPWNTSWSNKSLGQIWDNMTWMQWEKEINNYTGLIYTLIEQSQNQQEKNEQELLELDTWASLWNWFDISKWLWYIKIFIMIVGGLIGLRIVFTVLSIVNRVRQGYSPLSFQTRLPAQRG-PDRPEGIEEEGGERDRDRSGPLVNGFFAIIWVDLRNLCLFIYHRLRDLLLIVTRIVELLGR-RGWEILKYWWNLLQYWRQELKNSAISLLNATAIAVAEGTDRVIEVVQRIGRGILHIPVRIRQGLERALL

>V08DA_PRAp_2A5.5

MRVRGIRKNYQRLWKWG-TLLSMLLGLLMICNAKEQLWVTVYYGVPVWKEASTTLFCASDARAYSTEKHNVWATHACVPTDPDPQEVGLGNVTENFNMWKNNMVDQMHEDIISLWDQSLKPCVEITPLCVTLNCTDYRGNDNSTN------NSNETIVNKEEIKNCSFNITTSLSDKV-QERALFYMYDITPINN-----TTNTSYKLTRCNSSVITQACPKVSFEPIPIHYCAPAGFAILKCRQKNFNGTGPCTDVSTVQCTHGIRPVVSTQLLLNGSLAEGEVVIKSENFTDNSKNIIVQLKEAINITCTRPNNNTRKSINI--GPGRAIYATGEIIGDIRQAHCNISEEKWNKTLEQIVEKLQKQFG-NKSIIFAQPSGGDPETVMHSFNCGGEFFYCNTTRLFNSTWNDSS----TLSNGTGNNTITLPCRIKQVINMWQKVGKAMYAPPIAGQIRCSSNITGLILTRDGG----SNNMSNETFRPGGGNMKDNWRSELYKYKVVKIEPLGIAPTKAKRRVVQREKRAIG-IGAVFLGFLGAAGSTMGAASVTLTVQARQLLFGIVQQQNNLLRAIEAQQHLLQLTVWGIKQLQARVLAVERYLKDQQLLGIWGCSGKLICTTAVPWNTSWSNKSLEQIWDNMTWMQWEKEINNYTGLIYTLIEQSQNQQEKNEQELLELDTWASLWNWFDISKWLWYIKIFIMIVGGLIGLRIVFTVLSIVNRVRQGYSPLSFQTRLPAQRG-PDRPEGIEEEGGERDRDRSGPLVDGFFAIIWVDLRNLCLFIYHRLRDLLLIVTRIVELLGR-RGWEILKYWWNLLQYWRQELKNSAISLLNATAIAVAEGTDRVIEVVQRIGRGILHIPVRIRQGLERALL

>V06_MO_POVIp_2D8.7

MKVKGIRRNYQLLWRW----GIMLLGILRICNATEELWVTVYYGVPVWKEANTTLFCASDAKTYDTEAHNVWATHACVPTDPNPQEIVLGNVTENFNMWKNDMVEQMHEDIISLWDQSLKPCVKLTPLCVKLNCTDASTNTTNSTSTTNSTSGGEERMEKEEIKNCSFKITTSTREKV-KGSAFFYKTDVVSIAK------DNTSYKLIHCNTSVITQACPKVSFEPIPIHYCAPAGFAILKCNNKTFNGTGPCKNVSTVLCTHGIKPVVSTQLLLNGSLAEGEVIIRSENFTNNAKTIIVHLNESVVINCTRPNNNTRRSINM--GPGRAFYAT-GIIGDIRQAYCNISQGKWNYTLKQIVKKLREQFGENKTIVFNQSSGGDPEIVTHSFNCGGEFFYCNTTKLFNSTWNTSA----WHGYQEKNDTITLPCKIRQIINMWQEVGKAMYAPPIAGVIRCSSNITGLLLTRDGG----NESEGNEIFRPGGGDMRDNWRSELYKYKVVKIEPLGIAPTKAKRRVVQREKRAVG-IGALFLGFLGAAGSTMGAASITLTGQARLLLSGIVQQQNNLLRAIEAQQHLLQLTVWGIKQLQARILAVERYLKDQQLLGIWGCSGKLICTTAVPWNTSWSNKSMDNIWNNMTWMEWEREIDNYTGIIYNLLEDSQYQQEKNEKELLELDKWASLWNWFSITNWLWYIKIFIMIVGGLVGLRIVFAAFSIINRVRQGYSPLSFQTRFPAPRG-PDRPEGIEEEGGERDRDRSGRLVSGFLPLIWDDLRNLCLFSYHRLRDLLLIVTRIVELLGR-RGWEALKYWWNLLQYWGQELKNSAVSLLNAATIAVAEGTDRIIEVLQGTFRGILHIPTRIRQGLERALL

>V06MO_PRAp_1G3.2_b

MKVKGIRRNYQLLWRW----GIMLLGILMICNATEKLWVTVYYGVPVWKEANTTLFCASDAKAYDTEVHNVWATHACVPTDPNPQELLLENVTENFNMWKNDMVEQMHEDIISLWDQSLKPCVKLTPLCVKLNCTDANTSTTNSTSTTNSTSGGEERMEKEEIKNCSFNITTSTQEKV-KGSAFFYKIDIVPIGN----DTTNTSYKLLHCNTSVITQACPKVSFEPIPIHYCAPAGFAILKCNNEKFNGTGPCKNVSTVLCTHGIKPVVSTQLLLNGSLAEGEVMIRSENFTNNAKTIIVHLNESVVINCTRPNNNTRKSINM--GPGRAFYAT-GIIGDIRQAHCNISRTAWNNTLEKIVKKLREQFGKNKTIVFNQSSGGDPEIVTHSFNCGGEFFYCNTTKLFNSTWNTSM----WHGSNEQEENVTIPCKIRQIINMWQEVGKAMYAPPIAGVIRCSSEITGLLLTRDGG--NESESEGKEIFRPGGGDMRDNWRSELYKYKVVKIEPLGIAPTKAKRRVVQREKRAVG-IGALFLGFLGAAGSTMGAASIVLTGQARLLLSGIVQQQNNLLRAIEAQQHLLQLTVWGIKQLQARVLAVERYLKDQQLLGIWGCSGKLICTTAVPWNASWSNKSQNEIWNNMTWMEWEREIDNYTGIIYNLLEDSQYQQEKNEKELLELDKWASLWNWFSITNWLWYIRIFIMIVGGLVGLRIVFAVFSVINKVRQGYSPLSFQTRFPAPRG-PDRPEGIEEEGGERDRDRSGRLVNGFLPLIWDDLRSLCLFSYRRLRDLLLIATRIVELLGR-RGWEALKYWWNLLQYWGQELKNSAVSLLNAAAIAVAEGTDRIIEVLQRIFRAILHIPTRIRQGLERALL

>V06_MO_POVIp_1F2.7

MKVKGIRRNYQLLWRW----GIMLLGILRICNATEELWVTVYYGVPVWKEANTTLFCASDAKAYDKEVHNVWATHACVPTDPTPQEIVLGNVTENFNMWKNDMVEQMHEDIISLWDQSLKPCVKLTPLCVKLNCTDASTSTTNSTSTTNSTSGGEERMEKEEIKNCSFNITTSTQEKV-KGSAFFYKIDIVPIGN----DTTNTSYRLIHCNTSVITQACPKVSFEPIPIHYCAPAGFAILKCNNETFNGTGPCTNVSTVICTHGIKPVVSTQLLLNGSLAEGEVMIRSENFTNNAKTIIVHLNESVVINCTRPNNNTRKSINM--GPGRAFYAT-GIIGDIRQAHCNISRTAWNNTLKKIVEKLREQFGKNKTIVFNQSSGGDPEIVTHSFNCGGEFFYCNTTKLFNSTWNTSI----EHGSKDQEGNITLPCKIRQIINMWQEVGKAMYAPPIAGVIRCSSDITGLLLTRDGG--NESESEGKEIFRPGGGDMRDNWRSELYKYKVVKIEPLGIAPTKAKRRVVQREKRAVG-IGALFLGFLGAAGSTMGAASIMLTGQTRQLLSGIVQQQNNLLRAIEAQQHLLQLTVWGIKQLQARVLAVERYLKDQQLLGIWGCSGKLICTTDVPWNTSWSNKTLDKIWNNMTWMEWEREIDNYTGIIYNLLEDSQYQQEKNEKELLELDKWTSLWTWFSITNWLWYIRIFIMIVGGLVGLRIVFAVISIINKVRQGYSPLSFQTRFPAPRGRPDRPEGIEEEGGERDRDRSGRLVNGFLPLIWDDLRSLCLFSYRRLRDLLLIVTRIVELLGR-RGWEALKYWWNLLQYWGQELKNSAVSLLNAAAIAVAEGTDRIIEVLQRIFRAILHIPTRIRQGLERALL

>V06_MO_PRAp_1E2.6

MKVKGIRRNYQLLWKW----GIMLLGILMICNATEKLWVTVYYGVPVWKEANTTLFCASDARAYDKEVHNVWATHACVPTDPNPQEVVLGNVTENFNMWKNDMVEQMHEDIISLWDQSLKPCVKLTPLCVKLNCTNASTNTTNSTSTTNSTSDDKERMEKEEIKNCSFNITTSTQEKV-KGSAFFYKIDIVPMGN----DTTNTSYRLIHCNTSVITQACPKVSFEPIPIHYCAPAGFAILKCNNETFNGTGPCKNVSTVLCTHGIKPVVSTQLLLNGSLAEGEVMIRSENFTNNAKTIIVHLNEAVVINCTRPNNNTRRSINM--GPGRAFYAT-GIIGDIRQAHCNISRTAWNDTLKKIVKKLREQFG-NKTIVFNQSSGGDPEIVTHSFNCGGEFFYCNTTKLFSSTWNTST----WHESNDQGENITLPCKIRQIINMWQEVGKAMYAPPIAGVIRCSSNITGLLLTRDGG----NGSEGKEIFRPGGGDMRDNWRSELYKYKVVKIEPLGIAPTKAKRRVVQREKRAVG-IGALFLGFLGAAGSTMGAASIMLTGQTRLLLSGIVQQQNNLLRAIEAQQHLLQLTVWGIKQLQARVLAVERYLKDQQLLGIWGCSGKLICTTDVPWDTNWSNKTLDKIWNNMTWMEWEREIDNYTGIIYNLLEESQYQQEKNEKELLELDKWASLWNWFSITNWLWYIRIFIMIVGGLVGLRIVFTVLSIINRVRQGYSPLSFQTRFPAPRGRPDRPEEIEEEGGERDRDRSGRLVNGFLPLIWDDLRSLCLFSYRRLRDLILIVTRIVELLGR-RGWEALKYWWNLLQYWGQELKNSAVSLLNAAAIAVAEGTDRIIEVLQRIFRAILHIPTRIRQGLERALL

>V04EB_PRAp_1H5.6

MKIKETRKNYQHFWKW----GTMLLGMLLICSAAEQLWVTVYYGVPVWKEATTTLFCASDAKAYDTEAHNVWATHACVPTDPNPQEVVLGNVTENFNAWKNNMVEQMHEDVISLWDQSLKPCVKLTPLCVTLNCTDWKNPTNNTNTT-DTNATTVNREELGEIKNCSFNITTSMRDKMQKAYAPFYKLDVEPIDN------NNTSYRLISCNTSVITQACPKMSFEPIPIHYCAPAGFAILKCNNKTFNGKGPCTNVSTVQCTHGIKPVVSTQLLLNGSLAEKEVVIRSENFTDNAKVITVHLNDSVGINCTRPNNNTRKSIQI--GPGRAFFETGGIIGNIRQAHCNISQKQWNTTLGQIVVKLREQYGSNKTIVFNQSSGGDPEIVMHSFNCGGEFFYCNTTPLFNSTWNGTNG--TWKGTNGTNGTITIQCRIKQFINMWQGVGKAMYAPPISGIIRCSSNITGLLLTRDGG----NETSGTEIFRPAGGNMKDNWRSELYKYKVVKIEPLGVAPTKAKRRVVQREKRAVGTIGAMFLGFLGAAGSTMGAASMTLTVQARLLLSGIVQQQNNLLRAIEAQQHLLQLTVWGIKQLQARVLAVERFLRDQQLLGIWGCSGKLICTTTVPWNASWSNKSLDMIWHNMTWMQWEREIDNYTQLIYTLIEESQNQQEKNEQELLELDKWASLWNWFSITQWLWYIKISIMIVAGLVGLRIVFAVLSIVNRVRQGYSPLSFQTRLPTPRG-PDRPEETEEEGGDRDRDRSVRLVNGFLALIWDDLRSLCLFSYHHLRDLLLIAARIVELLGR-RGWEALKYWWNLLQYWIQELKNSAVSLLNATAIAVAEGTDRIIEVVQRACR------------ANRA--

>V04EB_POVIp_2A8.4

MKIKETRKNYQHFWKW----GTMLLGMLLICSAAEQLWVTVYYGVPVWKEATTTLFCASDAKAYDTEAHNVWATHACVPTDPNPQEVVLGNVTENFNAWKNNMVEQMHEDVISLWDQSLKPCVKLTPLCVTLNCTDWKNPTNNTNTT-DTNTTIVNREELGEIKNCSFNITTSMRDKMQKAYAPFYKLDVEPIDN------NNTSYRLISCNTSVITQACPKMSFEPIPIHYCAPAGFAILKCNNKTFNGKGPCTNVSTVQCTHGIKPVVSTQLLLNGSLAEKEVVIRSENFTDNAKVITVHLNDSVGINCTRPNNNTRKSIQI--GPGRAFFETGGIIGNIRQAHCNISQKHWNTTLGQIVVKLREQYGSNKTIVFNQSSGGDPEIVMHSFNCGGEFFYCNTTPLFNSTWNGTNG--TWKGTNGTNGTITIQCRIKQFINMWQGVGKAMYAPPISGIIRCSSNITGLLLTRDGG----NETSGTEIFRPAGGNMKDNWRSELYKYKVVKIEPVGVAPTKAKRRVVQREKRAVGTIGAMFLGFLGAAGSTMGAASMTLTVQARLLLSGIVQQQNNLLRAIEAQQHLLQLTVWGIKQLQARVLAVERFLRDQQLLGIWGCSGKLICTTTVPWNASWSNKSLDMIWHNMTWMQWEREIDNYTQLIYTLIEESQNQQEKNEQELLELDKWASLWNWFSITQWLWYIKIFIMIVAGLVGLRIVFAVLSIVNRVRQGYSPLSFQTRLPTPRG-PDRPEETEEEGGDRDRDRSVRLVNGFLALIWDDLRSLCLFSYHHLRDLLLIAARIVELLGR-RGWEALKYWWNLLQYWIQELKNSAVSLLNATAIAVAEGTDRIIEVVQRACR------------ANRA--

>V04EB_POVIp_1D1.1

MKIKETRKNYQHFWKW----GTMLLGMLLICSAAEQLWVTVYYGVPVWKEATTTLFCASDAKAYDTEAHNVWATHACVPTDPNPQEVVLGNVTENFNAWKNNMVEQMHEDVISLWDQSLKPCVKLTPLCVTLNCTDWKNPTNNTNTT-DTNATTVNREELGEIKNCSFNITTSMRDKMQKAYAPFYKLDVEPIDN------NNTSYRLISCNTSVITQACPKMSFEPIPIHYCAPAGFAILKCNNKTFNGKGPCTNVSTVQCTHGIKPVVSTQLLLNGSLAEKEVVIRSENFTDNAKVITVHLNDSVGINCTRPNNNTRKSIQI--GPGRAFFETGGIIGNIRQAHCNISQKQWNTTLGQIVVKLREQYGSNKTIVFNQSSGGDPEIVMHSFNCGGEFFYCNTTPLFNSTWNGTNG--TWKGTNGTNGTITIQCRIKQFINMWQGVGKAMYAPPISGIIRCSSYITGLLLTRDGG----NETSGTEIFRPAGGNMKDNWRSELYKYKVVKIEPLGVAPTKAKRRVVQREKRAVGTIGAMSLGFLGAAGSTMGAASMTLTVQARLLLSGIVQQQNNLLRAIEAQQHLLQLTVWGIKQLQARVLAVERFLRDQQLLGIWGCSGKLICTTTVPWNASWSNKSLDMIWHNMTWMQWEREIDNYTQLIYTLIEESQNQQEKNEQELLELDKWASLWNWFSITQWLWYIKIFIMIVAGLVGLRIVFAVLSIVNRVRQGYSPLSFQTRLPTPRG-PDRPEETEEEGGDRDRDRSVRLVNGFLALIWDDLRSLCLFSYHHLRDLLLIAARIVELLGR-RGWEALKYWWNLLQYWIQELKNSAVSLLNATAIAVAEGTDRIIEVVQRACR------------ANRA--

>V04EB_POVIp_1F8.7

MKIKETRKNYQHFWKW----GTMLLGMLLICSAAEQLWVTVYYGVPVWKEATTTLFCASDAKAYDTEAHNVWATHACVPTDPNPQEVVLGNVTENFNAWKNNMVEQMHEDVISLWDQSLKPCVKLTPLCVTLNCTDWKNPTNNTNTT-DTNATTTNREELGEIKNCSFNITTSMRDKMQKAYAPFYKLDVEPIDN------NNTSYRLISCNTSVITQACPKMSFEPIPIHYCAPAGFAILKCNNKTFNGKGPCTNVSTVQCTHGIKPVVSTQLLLNGSLAEKEVVIRSENFTDNAKVITVHLNDSVGINCTRPNNNTRKSIQI--GPGRAFFETGGIIGNIRQAHCNISQKQWNTTLGQIVVKLREQYGSNKTIVFNQSSGGDPEIVMHSFNCGGEFFYCNTTPLFNSTWNGTNG--TWKGTNGTNGTITIQCRIKQFINMWQGVGKAMYAPPISGIIRCSSNITGLLLTRDGG----NETSGTEIFRPAGGNMKDNWRSELYKYKVVKIEPLGVAPTKAKRRVVQREKRAVGTIGAMFLGFLGAAGSTMGAASMTLTVQARLLLSGIVQQQNNLLRAIEAQQHLLQLTVWGIKQLQARVLAVERFLRDQQLLGIWGCSGKLICTTTVPWNASWSNKSLDMIWHNMTWMQWEREIDNYTQLIYTLIEESQNQQEKNEQELLELDKWASLWNWFSITQWLWYIKIFIMIVAGLVGLRIVFAVLSIVNRVRQGYSPLSFQTRLPTPRG-PDRPEETEEEGGDRDRDRSVRLVNGFLALIWDDLRSLCLFSYHHLRDLLLIAARIVELLGR-RGWEALKYWWNLLQYWIQELKNSAVSLLNATAIAVAEGTDRIIEVVQRACR------------ANRA--

>V04EB_POVIp_1G4.5

MKIKETRKNYQHFWKW----GTMLLGMLLICSAAEQLWVTVYYGVPVWKEATTTLFCASDAKAYDTEAHNVWATHACVPTDPNPQEVVLGNVTENFNAWKNNMVEQMHEDVISLWDQSLKPCVKLTPLCVTLNCTDWKNPTNNTNTT-DTNATTTNREELGEIKNCSFNITTSMRDKMQKAYAPFYKLDVEPIDN------NNTSYRLISCNTSVITQACPKMSFEPIPIHYCAPAGFAILKCNNKTFNGKGPCTNVSTVQCTHGIKPVVSTQLLLNGSLAEKEVVIRSENFTDNAKVITVHLNDSVGINCTRPNNNTRKSIQI--GPGRAFFETGGIIGNIRQAHCNISQKQWNTTLGQIVVKLREQYGSNKTIVFNQSSGGDPEIVMHSFNCGGEFFYCNTTPLFNSTWNGTNG--TWKGTNGTNGTITIQCRIKQFINMWQGVGKAMYAPPISGIIRCSSNITGLLLTRDGG----NETSGTEIFRPAGGNMKDNWRSELYKYKVVKIEPLGVAPTKAKRRVVQREKRAVGTIGAMFLGFLGAAGSTMGAASMTLTVQARLLLSGIVQQQNNLLRAIEAQQHLLQLTVWGIKQLQARVLAVERFLRDQQLLGIWGCSGKLICTTTVPWNASWSNKSLDMIWHNMTWMQWEREIDNYTQLIYTLIEESQNQQEKNEQELLELDKWASLWNWFSITQWLWYIKIFIMIVAGLVGLRIVFAVLSIVNRVRQGYSPLSFQTRLPTPRG-PDRPEETEEEGGDRDRDRSVRLVNGFLALIWDDLRSLCLFSYHHLRDLLLIAARIVELLGR-RGWEALKYWWNLLQYWIQELKNSAVSLLNATAIAVAEGTDRIIEVVQRACR------------ANRA--

>V04EB_PRAp_2D9.5

MKIKETRKNYQHFWKW----GTMLLGMLLICSAAEQLWVTVYYGVPVWKEATTTLFCASDAKAYDTEAHNVWATHACVPTDPNPQEVVLGNVTENFNAWKNNMVEQMHEDVISLWDQSLKPCVKLTPLCVTLNCTDWKNPTNNTNTT-DTNATTVNREELGEIKNCSFNITTSMRDKMQKAYAPFYKLDVEPIDN------NNTSYRLISCNTSVITQACPKMSFEPIPIHYCAPAGFAILKCNNKTFNGKGPCTNVSTVQCTHGIKPVVSTQLLLNGSLAEKEVVIRSENFTDNAKVITVHLNDSVGINCTRPNNNTRKSIQI--GPGRAFFETGGIIGNIRQAHCNISQKQWNTTLGQIVVKLREQYGSNKTIVFNQSSGGDPEIVMHSFNCGGEFFYCNTTPLFNSTWNGTNG--TWKGTNGTNGTITIQCRIKQFINMWQGVGKAMYAPPISGIIRCSSNITGLLLTRDGG----NETSGTEIFRPAGGNMKDNWRSELYKYKVVKIEPLGVAPTKAKRRVVQREKRAVGTIGAMFLGFLGAAGSTMGAASMTLTVQARLLLSGIVQQQNNLLRAIEAQQHLLQLTVWGIKQLQARVLAVERFLRDQQLLGIWGCSGKLICTTTVPWNASWSNKSLDMIWHNMTWMQWEREIDNYTQLIYTLIEESQNQQEKNEQELLELDKWASLWNWFSITQWLWYIKTFIMIVAGLVGLRIVFAVLSIVNRVRQGYSPLSFQTRLPTPRG-PDRPEETEEEGGDRDRDRSVRLVNGFLALIWDDLRSLCLFSYHHLRDLLLIAARIVELLGR-RGWEALKYWWNLLQYWIQELKNSAVSLLNATAIAVAEGTDRIIEVVQRACR------------ANRA--

>V04EB_PRAp_2A10.7

MKIKETRKNYQHFWKW----GTMLLGMLLICSAAEQLWVTVYYGVPVWKEATTTLFCASDAKAYDTEAHNVWATHACVPTDPNPQEVVLGNVTENFNAWKNNMVEQMHEDVISLWDQSLKPCVKLTPLCVTLNCTDWKNPTNNTNTT-DTNATTVNREELGEIKNCSFNITTSMRDKMQKAYAPFYKLDVEPIDN------NNTSYRLISCNTSVITQACPKMSFEPIPIHYCAPAGFAILKCNNKTFNGKGPCTNVSTVQCTHGIKPVVSTQLLLNGSLAEKEVVIRSENFTDNAKVITVHLNDSVGINCTRPNNNTRKSIQI--GPGRAFFETGGIIGNIRQAHCNISQKQWNTTLGQIVVKLREQYGSNKTIVFNQSSGGDPEIVMHSFNCGGEFFYCNTTPLFNSTWNGTNG--TWKGTNGTNGTITIQCRIKQFINMWQGVGKAMYAPPISGIIRCSSNITGLLLTRDGG----NETSGTEIFRPAGGNMKDNWRSELYKYKVVKIEPLGVAPTKAKRRVVQREKRAVGTIGAMFLGFLGAAGSTMGAASMTLTVQARLLLSGIVQQQNNLLRAIEAQQHLLQLTVWGIKQLQARVLAVERFLRDQQLLGIWGCSGKLICTTTVPWNASWSNKSLDMIWHNMTWMQWEREIDNYTQLIYTLIEESQNQQEKNEQELLELDKWASLWNWFSITQWLWYIKIFIMIVAGLVGLRIVFAVLSIVNRVRQGYSPLSFQTRLPTPRG-PDRPEETEEEGGDRDRDRSVRLVNGFLALIWDDLRSLCLFSYHHLRDLLLIAARIVELLGR-RGWEALKYWWNLLQYWIQELKNSAVSLLNATAIAVAEGTDRIIEVVQRACR------------ANRA--

>V04EB_PRAp_2C10.1

MKIKETRKNYQHFWKW----GTMLLGMLLICSAAEQLWVTVYYGVPVWKEATTTLFCASDAKAYDTEAHNVWATHACVPTDPNPQEVVLGNVTENFNAWKNNMVEQMHEDVISLWDQSLKPCVKLTPLCVTLNCTDWKNPTNNTNTT-DTNATTVNREELGEIKNCSFNITTSMRDKMQKAYAPFYKLDVEPIDN------NNTSYRLISCNTSVITQACPKMSFEPIPIHYCAPAGFAILKCNNKTFNGKGPCTNVSTVQCTHGIKPVVSTQLLLNGSLAEKEVVIRSENFTDNAKVITVHLNDSVGINCTRPNNNTRKSIQI--GPGRAFFETGGIIGNIRQAHCNISQKQWNTTLGQIVVKLREQYGSNKTIVFNQSSGGDPEIVMHSFNCGGEFFYCNTTPLFNSTWNGTNG--TWKGTNGTNGTITIQCRIKQFINMWQGVGKAMYAPPISGIIRCSSNITGLLLTRDGG----NETSGTEIFRPAGGNMKDNWRSELYKYKVVKIEPLGVAPTKAKRRVVQREKRAVGTIGAMFLGFLGAAGSTMGAASMTLTVQARLLLSGIVQQQNNLLRAIEAQQHLLQLTVWGIKQLQARVLAVERFLRDQQLLGIWGCSGKLICTTTVPWNASWSNKSLDMIWHNMTWMQWEREIDNYTQLIYTLIEESQNQQEKNEQELLELDKWASLWNWFSITQWLWYIKIFIMIVAGLVGLRIVFAALSIVNRVRQGYSPLSFQTRLPTPRG-PDRPEETEEEGGDRDRDRSVRLVNGFLALIWDDLRSLCLFSYHHLRDLLLIAARIVELLGR-RGWEALKYWWNLLQYWIQELKNSAVSLLNATAIAVAEGTDRIIEVVQRACR------------ANRA--

>V02_RP_PRAp_2A12.3

MRVREIRRNSRQFWKW----GIMLLGMLMICSAVEQLWVTVYYGVPVWKDANTTLFCASDAKAYDTEVHNVWATHACVPTDPNPQEVLLENVTENFNMWKNNMVDQMHEDIISLWDQSLKPCVKLTPLCVTLDCTDLRNATNTTSN---DTSNGGGMPEGGEMKNCSFNITTSVRDKMQKTYALFYKLDVEPING------DNTSYRLISCNTSVITQACPKVTFEPIPIHYCAPAGFAILKCNDKKFNGTGQCRNVSTVQCTHGIKPVVSTQLLLNGSLAEEEVVIRSVNFTNNAKTIIVQLKEAVQINCTRPNNNTRKGIHI--GPGRAFYATGSIIGNIRQAYCNLSSTKWHNTLQQIVKKLREQFG-NKTIIFNQSSGGDPEIVMHSFNCGGEFFYCNSTQLFNSTWTNST-----EGLNTTEDPITLPCRIKQIINLWQEVGKAMYAPPISGIISCSSNITGLLLTRDGG--KNGSANNTEIFRPEGGDMRDNWRSELYKYKVVKIEPLGIAPTKAKRRVVQREKRAVG-LGAMFLGFLGAAGSTMGAAAITLTGQARQLLSGIVQQQNNLLRAIEAQQHMLQLTVWGIKQLQARVLAVERYLKDQQLLGIWGCSGKLICPTAVPWNTSWSNRSLEKIWDNMTWMEWEREIDNYTGLIYSLIEESQNQQEKNEQELLELDKWASLWNWFNITQWLWYIKIFIMIVGGLVGLRIVFAVFSIVNRVRQGYSPLSFQTLLPAPRG-PDRPEGIEEEGGERDRDRSGRLVTGFLALIWDDLRSLCLFSYHRLRDLLLIVARIVELLGR-RGWEALKYWWNLLKYWSQELKSSAISLLNVTAIAVAEGTDRIVEVLQRICRAFRNIPRRIRQGLERILI

>V02_POVIp_2D4.2

MRVREIRKSSRQFWKW----GIMLLGMLMICSAVEQLWVTVYYGVPVWKDANTTLFCASDAKAYDTEVHNVWATHACVPTDPNPQEVILENVTENFNMWKNNMVDQMHEDIISLWDQSLKPCVKLTPLCVTLDCTDLKNATNTTSN---GTSNGGGMPEGGEIKNCSFNITTSVRDKMQKTYALFYKLDVEPING------DNTSYRLISCNTSVITQACPKVTFEPIPIHYCAPAGFAILKCNDKKFNGTGQCRNVSTVQCTHGIKPVVSTQLLLNGSLAEEEVVIRSVNLTDNAKTIIVQLKEAVQINCTRPNNNTRKGIHI--GPGRAFYATGDIIGNIRQAHCNLSSTKWNNTLKQIVKKLREQFG-NKTIIFNQSSGGDPEIVMHTFNCGGEFFYCNSTKLFNSTWNSTE-----ELNTTEGPPITLPCRIKQIINLWQEVGKAMYAPPISGIISCSSNITGLLLTRDGG----NGSETNETFRPGGGNMRDNWRSELYKYKVVKIEPLGIAPTKAKRRVVQREKRAVG-LGAMFLGFLGAAGSTMGAAAITLTGQARQLLSGIVQQQNNLLRAIEAQQHMLQLTVWGIKQLQARVLAVERYLKDQQLLGIWGCSGKLICPTAVPWNTSWSNRSLEKIWNNMTWMEWEREIDNYTGLIYSLIEESQNQQEKNEQELLELDKWASLWNWFNITQWLWYIKIFIMIVGGLVGLRIVFAVFSIVNRVRQGYSPLSFQTLLPAPRG-PDRPEGIEEEGGERDRDRSGRLVTGFLALIWDDLRSLCLFSYHRLRDLLLIVARIVELLGR-RGWEALKYWWNLLKYWSQELKSSAISLLNVTAIAVAEGTDRIVEVLQRTCRAFLNIPRRIRQGLERILI

>V02RP_POVIp_1A5.8_2

MRVREIRKSSRQFWKW----GIMLLGMLMICSAVEQLWVTVYYGVPVWKDANTTLFCASDAKAYDTEVHNVWATHACVPTDPNPQEVILENVTENFNMWKNNMVDQMHEDIISLWDQSLKPCVKLTPLCVTLDCTDLKNATNTTSN---GTSNGGGMPEGGEMKNCSFNITTSVRDKMQKTYALFYKLDVEPING------DNTSYRLISCNTSVITQACPKVTFEPIPIHYCAPAGFAILKCNDKKFNGTGQCRNVSTVQCTHGIKPVVSTQLLLNGSLAEEEVVIRSVNLTDNAKTIIVQLKEAVQINCTRPNNNTRKGIHI--GPGRAFYATGDIIGNIRQAHCNLSSTKWNNTLKQIVKKLREQFG-NKTIIFNQSSGGDPEIVMHTFNCGGEFFYCNSTKLFNSTWNSTE-----ELNTTEGPPITLPCRIKQIINLWQEVGKAMYAPPISGIISCSSNITGLLLTRDGG----NGSETNETFRPGGGNMRDNWRSELYKYKVVKIEPLGIAPTKAKRRVVQREKRAVG-LGAMFLGFLGAAGSTMGAAAITLTGQARQLLSGIVQQQNNLLRAIEAQQHMLQLTVWGIKQLQARVLAVERYLKDQQLLGIWGCSGKLICPTAVPWNTSWSNRSLEKIWNNMTWMEWEREIDNYTGLIYSLIEESQNQQEKNEQELLELDKWASLWNWFNITQWLWYIKIFIMIVGGLVGLRIVFAVFSIVNRVRQGYSPLSFQTLLPAPRG-PDRPEGIEEEGGERDRNRSGRLVTGFLALIWDDLRSLCLFSYHRLRDLLLIVARIVELLGR-RGWEALKYWWNLLKYWSQELKSSAISLLNVTAIAVAEGTDRIVEVLQRTCRAFLNIPRRIRQGLERILI

>V09JF_PRAp-2A11.4

MKAKEIRKNCQRLWRW----GILLLGMLMICSATEKLWVTVYYGVPVWKEANTTLFCASDAKAYDTEAHNVWATHACVPTDPNPQEVVLANVTENFNMWKNDMVEQMHQDIISLWDQSLKPCVKLTPLCVTLNCTNLNTNNTNNSSGETNNNANNSSGDYGEIKNCSFNITTPIRNKVQKEYALFNRLDIVPIDD------DNASYTLISCNTSVITQACPKISFDPIPIHYCAPAGFAILKCNNKTFNGTGPCTNVSTVQCTHGIRPVVSTQLLLNGSLAEKETVIRSENFTNNAKTIIVQLNEPIPINCTRPNNNTRKSIHI--APGRAFYATGTIIGNIRQAHCNISNTTWQNALKNVVAKLREQFG-NKTIVFNQSSGGDPEITMHTFNCGGEFFYCDTTRLFNSTWGFNDT---SDRANSTNSNITLPCRIKQIINRWQEVGRAMYAPPIRGQIRCSSNITGLLLTRDGGNETEENKTNTETFRPGGGNMKDNWRSELYKYKVVKIEPLGVAPTKAKRRVVQREKRAVG-LGAMFLGFLGAAGSTMGAASITLTVQARQLLSGIVQQQNNLLRAIEAQQHMLQLTVWGIKQLQARVLAVERYLKDQQLLGLWGCSGKLICTTAVPWNTSWSNKSQTEIWNNMTWMEWEREINNYTKVIYTLIEESQNQQEKNEQDLLELDTWAGLWNWFDITNWLWYIKIFIMIVGGLIGLRIVFTVLSIVNRVRQGYSPLSLQTLFPGPRG-PDRPEGIEEGGGERDRNGSRTLVHGFLALVWVDLRSLCLFSYHRLRDLLLIVTRTVELLGR-RGWEALKYWWNILQYWSQELRSSAVSLLNATAIAVAEGTDRIIEVVQRACRAILHIPRRIRQGAERLLL

>V09JF_POVIp-1D4.2

MKAKEIRKNCQRLWRW----GVLLLGMLMICSATEKLWVTVYYGVPVWKEANTTLFCASDAKAYDTEAHNVWATHACVPTDPNPQEVVLANVTENFNMWKNDMVEQMHQDIISLWDQSLKPCVKLTPLCVILNCTNLNTNNTNN-------SSGDYGEIKGEIKNCSFNITTPIRNKVQKEYALFNRLDIVPIDD------DNASYTLISCNTSVITQACPKISFEPIPIHYCAPAGFAILKCNNKTFNGTGPCTNVSTVQCTHGIRPVVSTQLLLNGSLAEKETVIRSENFTNNAKTIIVQLNESIPIDCIRPNNNTRKSIHI--APGRAFYATGTIIGDIRQAYCNISNTTWQNALKKIVTKLREQFG-NKTIVFNKSSGGDPEITMHTFNCGGEFFYCDTTGLFNSTWGFNDT---SDWAKSTDSNITLPCRIKQIINRWQEVGRAMYAPPIRGQIRCSSNITGLLLTRDGGNETEENKTTTETFRPGGGNMKDNWRSELYKYKVVKIEPLGVAPTQAKRRVVQREKRAVG-LGAMFLGFLGAAGSTMGAASITLTVQARQLLSGIVQQQNNLLRAIEAQQHMLQLTVWGIKQLQARVLAVERYLKDQQLLGLWGCSGKLICTTAVPWNTSWSNKSQKEIWDNMTWMEWEREINNYTKVIYTLIEESQNQQEKNEQELLALDTWASLWNWFDITKWLWYIKIFIMIVGGLIGLRIVFTVLSIVNRVRQGYSPLSLQTLFPGPRG-PDRPEGIEEGGGERDRNGSRTLVHGFLALVWVDLRSLCLFSYHRLRDLLLIVTRTVELLGR-RGWEALKYWWNILQYWSQELRNSAVSLLNATAIAVAEGTDRIIEVVQRACRAILHIPRRIRQGAERLLL

>V09F_PRAp-2F9.2

MKAKEIRKNCQRLWRW----GILLLGMLMICSATEKLWVTVYYGVPVWKEANTTLFCASDAKAYDTEAHNVWATHACVPTDPNPQEVVLANVTENFNMWKNDMVEQMHQDIISLWDQSLKPCVKLTPLCVTLNCTNLNTNNTNNSSR-ETSNANNSSGDYGEIKNCSFNITTPIRNKVQKEYALFNRLDIVPIDD------DNASYTLISCNTSVITQACPKISFDPIPIHYCAPAGFAILKCNNKTFNGTGPCTNVSTVQCTHGIRPVVSTQLLLNGSLAEKEIIIRSENFTNNAKTIIVQLNESIPINCIRPNNNTRKSIHI--APGRAFYATGTIIGNIRQAHCNISNTTWQNALKKIVTKLREQFG-NKIIVFNQSSGGDPEITMHTFNCGGEFFYCDTTRLFNSTWEFNDT---SDQANSTNINITLPCRIKQIINRWQEVGRAMYAPPIRGQIRCSSNITGLLLTRDGGNETEENKTNTETFRPGGGNMKDNWRSELYKYKVVKIEPLGVAPTQAKRRVVQREKRAVG-LGAMFLGFLGAAGSTMGAASITLTVQARQLLSGIVQQQNNLLRAIEAQQHMLQLTVWGIKQLQARVLAVERYLKDQQLLGLWGCSGKLICTTAVPWNTSWSNKSQTEIWNNMTWMEWEREINNYTKVIYTLIEESQNQQEKNEQELLALDTWASLWNWFDITKWLWYIKIFITIVGGLIGLRIVFTVLSIVNRVRQGYSPLSLQTLFPGPRG-PDRPEGIEEGGGERDRNGSRTLVHGFLALVWVDLRSLCLFSYHRLRDLLLIVTRTVELLGR-RGWEALKYWWNILQYWSQELKNSAISLLNATAIAVAEGTDRIIEVVQRACRAILHIPRRIRQGAERILL

>V03NP_POVIp-2D5.10.2

MRVRGIRKNYQHWWGWS-TMLTMLLGMLMICKAAEQLWVTIYYGVPVWKEATTTLFCASDAKAYDTEMHNVWATHACVPTDPNPQEVVLENVTENFNMWENNMVEQMHEDIISLWDQSLKPCVKLTPLCVTLNCTNANISSNTSN------IYNSSIYKPGDIKNCSFNITTHIRDKVKKEYALFYALDVIPIDPTVGNKTTTTDYRLISCNTSVVTQACPKVSFEPIPIHYCAPAGFAILKCNDKMFNGTGPCKNVSTIQCTHGIRPVVSTQLLLNGSLAEEDIVIRSKNFTDNAKTIIVQLNKTVTINCTRPSNNTRKSIHI--APGRAFYATGNIIGDIRQAHCNLSRKDWNSTLRLIANKLREQFG-NKTIVFNSSSGGDPEIVMHSFNCRGEFFYCNTTPLFNSTWYSNN----TQEGMEGNSIITLQCRIKQIVNMWQEVGKAMYAPPIKGQIRCSSNITGLLLTRDGG----NTTNRPEVFRPGGGNMKDNWRSELYKYKVVKLEPLGVAPTKAKRRVVQREKRAVG-IGALFLGFLGAAGSTMGAASMTLTVQARQLLSGIVQQQNNLLMAIEAQQHMLQLTVWGIKQLQARVLAVESYLKDQQLLGIWGCSGKLICTTTVPWNTSWSNKTYNEIWDNMTWMQWEKEIDNHTSLIYTLIEESQNQQEKNELDLLALDKWANLWSWFNITNWLWYIKIFIMIVGGLVGLRIVFTVLSIVNRVRQGYSPLSFQTHLPARRG-PDRPEEIEEEGGERDRGRSEPLVTGFLTLIWVDLRSLCLFSYHRLRDLLLIAARIVELLGR-RGWEILKYWWNLLQYWSQELKNSAVSLLNATAIAVAEGTDRVIEIVQGIGRAILHIPARIRQGLERVLL

